# A novel pan class-I glucose transporter inhibitor DRB18 exhibits synergistic effects with paclitaxel *in vitro* and *in vivo* against human non-small cell lung cancer

**DOI:** 10.1101/2024.12.15.628558

**Authors:** Pratik Shriwas, Lindsey Bachmann, Ryan Ward, Subhodip Adhicary, Corinne M. Nielsen, Yunsheng Li, Haiyun Zhang, Jingwen Song, Dennis Roberts, Stephen Bergmeier, Xiaozhuo Chen

## Abstract

**Purpose:** Cancer cells depend on glucose for biomass synthesis, cell proliferation, and drug resistance. Glucose transporter (GLUT) transcripts as well as proteins are upregulated in human lungs and other cancers and are negatively correlated with patient survival, particularly GLUT1 and GLUT3. Thus, inhibiting GLUT function has been an attractive anticancer strategy. We previously characterized WZB117 and DRB18, first- and second-generation pan-class I GLUT inhibitors, respectively. DRB18 strongly inhibits glucose transport mediated by GLUT1-4 in non-small lung cancer (NSCLC) A549 cells *in vitro* and *in vivo*. Here, we report DRB18 as a more stable and potent anticancer compound, compared to WZB117.

**Methods:** Immunohistochemistry analysis was performed in Lung adenocarcinoma (LUAD) tissue array to investigate GLUT1 and GLUT3 protein expression between normal, lower and higher stage LUAD patients. Bioinformatics analysis was performed to examine additive effect of GLUT3 to GLUT1 mediated prognosis in LUAD. Glucose uptake and resazurin dye-based proliferation assays were used to determine glucose uptake inhibitory and cell proliferation inhibitory against panel of human cancer cell lines A549, Panc1 and Hela. DRB18 potency was tested against the presence of extracellular nutrients glucose, glutamine and ATP. Synergism between DRB18 and clinically approved anticancer drugs was tested against cancer cells. DRB18 and advanced NSCLC drug Paclitaxel were tested for synergy *in vitro* and *in vivo*.

**Results:** GLUT1/3 combination exhibited higher hazard ratio than either GLUT1 and GLUT3 alone in many cancer types including LUAD. DRB18 reduced glucose uptake in NSCLC A549, pancreatic Panc1, and cervical Hela cancer cells with varied but strong anticancer potencies in the presence or absence of extracellular nutrients such as ATP and glucose. Combined with different clinical and pre-clinical anticancer compounds such as V9302, CB839, Sutent, Brigatinib, DRB18 significantly increased death of A549 and Panc1 cells. Noteworthy, DRB18 exhibited strong anticancer synergy with paclitaxel, an approved chemo drug for NSCLC, drastically reducing cancer cell proliferation *in vitro* and growth of A549 tumors grafted on the flank of nude mice without significant side effects, compared to single drug treatments.

**Conclusions:** Collectively, our results demonstrate anticancer efficacy of pan class-I GLUT inhibitor DRB18 in combination with paclitaxel, providing a potentially more efficacious therapeutic strategy for treating advanced NSCLC and other cancers.

## Background

Lung cancer is the second most common cancer among men and women in the United States [1], with non-small cell lung cancer (NSCLC) contributing to more than 80% of all lung cancer cases [2]. Currently, major therapeutic strategies for NSCLC include chemo or targeted therapy and immune therapy, with chemotherapy remaining the gold standard in NSCLC treatment [3, 4]. Despite these different therapies, the five-year survival rate of NSCLC remains under 30% [2]. Major issues in lung cancer therapy include resistance developed to therapy, and side-effects at higher dosages of drugs [5, 6]. It has been established that combinatorial drug therapies are more effective in treating different types of cancers and induce less drug resistance than single drug therapies [7, 8]. It has been reported that drug resistance in cancer, including NSCLC, are closely associated with glucose transport and glucose metabolism [9, 10].

Glucose uptake and anerobic glycolysis are drastically upregulated in different cancer types for the synthesis of biomass, energy production, and cell proliferation. This phenomenon is known as the Warburg effect [11–13]. Glucose uptake is mediated by a group of plasma membrane-spanning protein channels called glucose transporters (GLUTs) [14–16]. GLUT1 is the main GLUT in most human cells and most cancer cells, internalizing glucose to satisfy the energetic requirements of the cell [17, 18]. It is well known that GLUTs, particularly class I GLUTs (GLUT1-4), are either inducible or co-expressed with GLUT1 in different types of cancer [19, 20]. Further studies show that these GLUTs possess unique, but also partially redundant, functions with each other and frequently compensate in case of single GLUT inhibition, making successful single-GLUT or specifically GLUT1 targeting more challenging.

Consequently, targeting glucose transport and metabolism, via inhibiting GLUTs, becomes an attractive anticancer strategy [21–23]. Recently, many different GLUT inhibitors have been developed and tested [24–27] in pre-clinical models. WZB117, a first-generation lead inhibitor synthesized in our labs [28–30], has shown anticancer activity across several cancers via reducing drug resistance and epithelial mesenchymal transition (EMT) of cancer cells [29–31] as well as exhibiting additive anticancer effects, *in vitro* and *in vivo*, in combination with FDA-approved drugs [32].

We recently showed that knocking out the *GLUT1* gene in NSCLC A549 cells did not reduce tumor growth in Nu/J mice [38], suggesting that other class-I GLUTs might have compensated for the lost GLUT1 activity. In contrast, simultaneous inhibition of multiple GLUTs, by second-generation lead GLUT inhibitor DRB18 [38], appears to be a far more effective strategy to block glucose transport and glucose metabolism, ultimately leading to increased cancer cell death. Demonstrated by a molecular docking study, DRB18 binds human GLUT1-4 in an outward open conformation, exhibiting glucose uptake inhibitory IC_50_s varying from 0.9 µM to 9 µM among human GLUT1-4. DRB18 reduced growth in A549 xenograft tumors at 10 mg/kg body weight, when administered intraperitoneally daily for five weeks [38].

Observing the targeting of pan Class I GLUTs and the improved anticancer efficacy of DRB18, we hypothesized that DRB18 could potentially be used in combination with approved anticancer drugs to achieve synergistic anticancer efficacy. During combinatorial therapy, we lowered the dosages of DRB18 and paired anticancer agents and achieved greater tumor inhibitory activities, with lower side effects of each compound. These results highlight the potential of DRB18, alone or combined with approved anticancer drugs, to enhance therapeutic efficacy without significant side effects

## Methods

### Compound inhibitors and other chemicals

Compounds WZB117 and DRB18 were synthesized as previously reported [28,35]. Compound solutions were freshly prepared by dissolving the compounds in dimethyl sulfoxide (DMSO) before each experiment. Adenosine 5-triphosphate disodium salt hydrate (A2383), D-(+)-Glucose (G7021), L-glutamine solution 200 mM (59202-C), Paclitaxel (T7402), Sunitinib malate (PZ0012), and Resazurin sodium salt (R7017) were purchased from Sigma Aldrich. Brigatinib (19778) and Trametinib (16292) were purchased from Cayman chemicals. V9302 (HY-112683) and CB839 (HY-12248) were purchased from MedChemExpress. Dulbecco’s Modified eagle Medium (DMEM) high glucose (ATCC-30-2002), Eagle’s Minimum Essential Medium (EMEM-ATCC 30-2003), RPMI (ATCC-30-2001), Fetal bovine serum (ATCC-30-2020), and Penicillin-Streptomycin Solution (ATCC-30-2300) were purchased from American Type Culture Collection (ATCC). DMEM (no glucose, no glutamine) was purchased from Thermo Fisher Scientific (A1443001). Deoxy-D-glucose, 2-[1,2-3H(N)], 250 µCi (37 MBq) was purchased from Perkin Elmer (NET549250UC). Stock concentrations of compounds were prepared by dissolving compounds in DMSO or Ethanol. The working solutions of the compounds were prepared by diluting stock solutions in appropriate cell culture media.

## Cell lines and experimental controls

All cell lines were purchased from ATCC and were grown in growth media supplemented with 10% FBS and 1% penicillin/streptomycin (Pen/strep). Dulbecco’s Modified Eagle Medium (DMEM) was growth media for A549, H1299, and Panc1 cells, EMEM for HOP92 and Hela cells while INS-1 cells were cultured in RPMI-1640 [39]. All the cells were grown in a cell culture incubator with 5% CO_2_ at 37^0^C. Mock (DMSO/Ethanol) treated samples served as negative controls.

## Glucose uptake assay in cancer cells and in specific GLUT-expressing cell lines

Glucose uptake inhibitory activity of DRB18 was analyzed using radioactive 2-deoxy-d-[^3^H] glucose as previously described (26,36). For initial seeding density of 40000 cells per well were used for all cell lines but 100,000 cells for INS-1 cells.

## Cell proliferation assays

Cell proliferation assays were performed using resazurin as a fluorescent dye as previously described [38]. For cell viability assays for extracellular nutrients, an appropriate amount of glucose, glutamine or ATP was added to the DMEM (no glucose/no glutamine) supplemented with 10% FBS and 1% pen/strep to prepare assay media. The cells were then treated with different concentrations of compounds or nutrients for 24, 48, or 72 hours. After treatment, the media was removed from cells and cell viability was measured using resazurin dye.

## Intracellular ATP assay

25,000 A549, Panc1 or Hela cancer cells were seeded in each well of 96 well black plate. Intracellular ATP levels in the cancer cells were measured as described previously [38].

## WZB117/DRB18 stability assay

WZB117 and DRB18 were pre-incubated at 37°C in 1 ml cell culture media at 30 µM concentration for different times (0-72 hours). A549 cells were seeded at 5,000 cells per well in a 96-well plate. Cells were then treated for 24 hours with the WZB117 or DRB18 pre-incubated solution. For the control, 0.1% DMSO was used to treat the cells for 24 hours. The cell viability was measured by a resazurin assay as described above.

## DRB18 synergism assay

5,000 A549/Panc1 cells were seeded in each well of 96-well plates. Cells were treated with appropriate concentrations of compounds for 24 or 48 hours. Cell viability was measured using resazurin dye as described above. Synergy score analysis was performed using SynergyFinder ver3.0 (https://synergyfinder.fimm.fi). An HSA model was used to determine synergism based on synergy scores, which were interpreted using the following baselines: score < -10: antagonistic, score between -10 and 10: additive; score >10: synergistic.

## Bioinformatics analysis

mRNA/protein correlation and copy number variation analysis of human GLUTs was performed using data collected from cbioportal (https://www.cbioportal.org/).

## TCGA-LUAD RNA-seq and Clinical Data

Primary RNA-seq expression (HTSeq-counts or FPKM), clinical metadata, mutation (MAF) files, and copy number alteration (CNA) files for lung adenocarcinoma (TCGA-LUAD) were downloaded using the TCGAbiolinks R package. Data were harmonized to the GRCh38 genome build. Clinical information included survival time, vital status, tumor stage, and nodal metastasis status. Raw counts were normalized using: edgeR::cpm(count_matrix, log = TRUE, prior.count = 1). This generated log2(CPM+1) normalized expression used for Stage-wise GLUT1/GLUT3 comparisons, Survival analysis, DEG analysis, Volcano plot, Heatmaps and clustering. Batch effects were assessed and corrected using limma::removeBatchEffect when required (not shown in Fig.s). Definition of GLUT1/GLUT3 Groups. To study cooperative effects of GLUT1 and GLUT3, samples were classified as: High-High (HH): Both GLUT1 and GLUT3 above median Or Low-Low (LL): Both GLUT1 and GLUT3 below median

## GSE68465 Microarray LUAD Dataset

The independent validation cohort GSE68465 (NSCLC microarray dataset with survival data) was downloaded using GEOquery. RMA-normalized values were used for expression analysis. GLUT1/GLUT3 high–high vs low–low groups were defined using median split for survival analysis, Cox regression, and GSEA. Microarray data were background-corrected, normalized (RMA), and probe-to-gene collapsed using the WGCNA collapse Rows function. GLUT1 and GLUT3 groups were created in a similar way as defined above.

## Statistical Analysis

### Differential Expression (DEG) Analysis

DEGs between GLUT1/GLUT3 HH vs LL groups were identified using:

design <- model.matrix(∼ group)
fit <- limma::lmFit(norm_expr, design)
fit2 <- limma::eBayes(fit)
deg <- limma::topTable(fit2, coef = 2, number = Inf)
Significance threshold: |log2FC| > 1 & FDR < 0.05.
**Volcano plots** were generated using EnhancedVolcano.

## Gene Set Enrichment Analysis (GSEA)

GSEA was performed using clusterProfiler with MSigDB Hallmark and KEGG gene sets. Pre-ranked GSEA with log2FC was used:

clusterProfiler::GSEA(pre_ranked_vector, TERM2GENE = msigdb_set)

Enriched pathways included cell cycle, glycolysis, hypoxia, MYC targets, etc. (Fig 2C).

Survival Analysis

Overall survival (OS) was modeled using:

- Kaplan–Meier Survival Curves (survminer::ggsurvplot)
- Cox Proportional Hazards Regression (survival::coxph)
- Forest Plots for hazard ratios (forest model package)

Analyses performed:

1. GLUT1 high vs low
2. GLUT3 high vs low
3. Combined GLUT1/3 HH vs LL
4. Subtype-specific survival (e.g., TCGA-PAAD, STAD, ACC, SARC, MESO)

**Cox models** included covariates:

coxph(Surv(time, status) ∼ GLUT1 + GLUT3 + age + stage, data)

## Heatmap and Clustering

The top 50 DEGs were extracted (upregulated and downregulated separately). Normalized expression values were Z-scaled and visualized using pheatmap with:

- hierarchical clustering (Euclidean)
- annotation for HH vs LL groups

## Immunohistochemistry study of human LUAD array

Fixed, paraffin-embedded human lung tissue samples (LC486, LC1531, HuCAT232, HuFPT178) (https://www.tissuearray.com/) included Stage I-IV lung adenocarcinoma and tumor/cancer adjacent controls (Supplementary Table 1-4). These clinical data are provided directly from suppliers. The data presented here are in compliance with their policy. To prepare tissue sections for antibody staining, samples were deparaffinized in xylenes, rehydrated in ethanol series, and subject to antigen retrieval using 10 mM sodium citrate + 0.05% Tween-20 at sub-boiling temperature (microwave oven), incubated in 3% hydrogen peroxide to quench endogenous peroxidase, and blocked in 5% donkey serum in 0.1% Tween-20 buffer. Rabbit anti-GLUT1/SLC2A1 (1:200, Proteintech 21829-1-AP) or Rabbit anti-GLUT3/SLC2A3 (1:50, AbClonal A8150) primary antibodies were applied in blocking reagent overnight at 4°C. HRP-conjugated donkey anti-rabbit (1:100, Cell Signaling 7074P2) secondary antibody was applied in blocking reagent for 2 hours at room temperature. For chromogen precipitation, 1% DAB chromogen (Electron Microscopy Sciences) + 0.3% hydrogen peroxide was applied for ∼5 min before reaction was stopped and tissue sections were mounted with Permount (Fisher). Patient characteristics are given in the Supplementary Table 1-4.

## Animal study

The animal experiments were performed in accordance with guidelines for animal care by the NIH and IACUC protocol approved by Ohio University.

4 weeks old male *Nu/J* mice were purchased from Jackson Laboratories and were put on Irradiated Teklad Global 19% rodent (protein) diet from Harlan Laboratories. The protocols for subcutaneous injection of tumors (tumor generation), intraperitoneal administration of compounds/vehicles, weekly tumor measurements, body weight/food intake measurements and animal euthanasia were performed as previously described [28, 38]. Briefly, mice were allowed to acclimatize for one week after arrival. They were injected with 5x10^6^ A549 cells subcutaneously on the flank. The tumors became palpable after 3 days at ∼100 mm^3^ volume. Mice were divided into 4 groups: Group I: control group (*n* = 8) treated with PBS/DMSO (1:1, v/v); Group II: 8 mg/kg (body weight) DRB18 treatment group (*n* = 7); Group III: 8 mg/kg (body weight) paclitaxel treatment group (*n* = 8); Group IV 8 mg/kg DRB18 + 8mg/kg paclitaxel treatment group (*n* = 8). Tumor volume, body weight and food intake were measured weekly for all the groups. The mice were euthanized after 5 weeks of treatment and final tumor volume and body weight were measured. All animals were anesthetized using isoflurane inhalation to ensure they were unconscious and insensitive to pain. The mice were euthanized using cervical dislocation. Isoflurane is a standard anesthetic that was selected due to fast action and reliability in terms of safety. Tumor volume was calculated by the formula; V= LW^2^/2 where L=Length, W= Width.

## Data processing and statistical analysis

For all the cell and molecular studies, each experimental condition was performed in triplicates or quadruplets and the experiment was repeated at least once. For animal studies, the data is represented as mean ± standard error of the means (SEM). Other data is reported as mean ± standard deviation (STD) and analyzed using Student’s two-tailed *t*-test. *P* ≤ 0.05 was considered significant.

## Data Availability and data sharing

The datasets used and/or analysed during the current study are available from the corresponding author on reasonable request.

## Results

### 3-1. GLUT1 and GLUT3 together contribute to poor survival in lung adenocarcinoma

*GLUT1* and *GLUT3* mRNA expression levels were found to be higher in tumors of human patients, compared to benign samples (Fig. S1A-B). Thus, we investigated the expression of GLUT1 and GLUT3 in human lung adenocarcinoma (LUAD) cancer tissue microarray of LUAD patients of different stages (Supplementary table T1-T4, Supplementary data 1). We found that GLUT1 protein expression was increased in Stages I & II tumors compared to normal lung tissue (P<0.0001), and GLUT1 was further upregulated in tumor samples of higher stage patients compared to lower stage (P=0.0064) (Fig. 1A-B). Similarly, we found that GLUT3 protein was increased in healthy tissue adjacent to tumor tissues of Stage I & II (P=0.0014), but it was not further increased in higher stage samples (P:NS) (Fig. 1C). In addition, we found that the expressions of GLUT1 and GLUT3 were higher in grade 3 tumors compared to grade 2 (CPTAC, APOLLO 2022 LUAD cohort n=87) (Fig. S1C-D, Supplementary data 2). Subsequently, GLUT1 and GLUT3 were significantly positively correlated (R=0.3458; P<0.0001) at the mRNA level (Fig. 1D), indicating coordinated transcriptional regulation and potential co-dependency in LUAD biology. We found that *GLUT1* mRNA expression was increased from lower (T1-T2) compared to higher (T3-T4) stages as well as based on nodal metastasis status (lower in patients without metastasis, N0 compared to those with metastasis N1/N2/N3). However, *GLUT3* mRNA was not increased significantly in higher stage patients as well as patients with nodal metastasis (Fig. S1E-H, Supplementary data 3). We then combined *GLUT1* and *GLUT3* mRNA score to investigate the combined score changes based on tumor stage and found that combined score was significantly higher (P=0.024) in higher stage patients indicating that they contribute together to disease progression (Fig. 1E, Supplementary data 4). Next, we found that LUAD patients from TCGA cohort which had higher *GLUT1* and *GLUT3* mRNA had poorer survival compared to those with lower expression (Fig. 1F). Further, we found that expressions of GLUT1 and GLUT3 were positively correlated in TCGA-LUAD (mRNA) (Fig. 1G, Supplementary data 2) as well as at protein level (Fig. S2A, Supplementary data 2). However, *GLUT4* was found to be negatively correlated with both *GLUT1* and *GLUT3* mRNA levels in the TCGA-LUAD cohort (Fig. S2B-C, Supplementary data 2). In the CCLE database (Broad Institute, 2019) GLUT1 and GLUT3 were found to be positively correlated at mRNA and proteins levels in overall cancer types (Fig. S2D-E) as well as lung cancer only (Fig. S2F-G, Supplementary data 2). We further found that *GLUT3* mRNA had the highest mutations (8%) among *GLUT1-4* in TCGA-LUAD cohort using OncoPrint tool in cbioportal (Fig. S3A). Copy number alterations in *GLUT3* mRNA expression levels in TCGA-LUAD cohort were increased significantly in patients with gain/amplifications and diploid compared to patients with deletions (Fig. S3B, Supplementary data 3). Protein-protein interaction study using genemania (https://genemania.org/) identified 20 major interacting partners of GLUT1 and GLUT3 (Fig. S3C). Similarly, string database analysis revealed p53 as a major interacting partner of GLUT1 and CREB1 for GLUT3 (Fig. S3D). Gene Ontology (GO) analysis using shinyGOv0.741 (http://bioinformatics.sdstate.edu/go74/) using the top interacting partners for GLUT1 and GLUT3 revealed NADH regeneration, canonical glycolysis, and glucose catabolism to pyruvate as major GO: biological processes; glucokinase, hexokinase, and fructokinase were major GO: molecular functions (Fig. S3E-F). Once we confirmed the variations in mRNA or protein expression for GLUT1 and GLUT3 as well their interacting partners, we next investigated survival in TCGA-LUAD patients based on expression of GLUT1-13 individually (except for GLUT2 where we could not divide the patients due to lack of data) (Fig. S4, Supplementary table 4). We wanted to investigate whether GLUT1 and GLUT3 contribute together to poor survival in LUAD. So, we divided TCGA-LUAD patients into GLUT1/3 high-high vs low-low groups based on whether the patient had higher expression of both mRNAs and found that high-high group had poorer survival (P=0.0064) compared to low-low group. (Fig. 1E, Supplementary data 4). Multivariate cox modelling regression was performed to validate survival result from Kaplan Meier analysis. We found that the hazard ratio in GLUT1/3 combo group (HR = 1.72, 95% CI: 1.18–2.51) for high-high vs low-low patients was higher than either GLUT1 (HR = 1.26, 95% CI: 1.14–1.40) or GLUT3 (HR = 1.04, 95% CI: 0.93–1.16) alone suggesting additive effect of GLUT3 to GLUT1 (Fig. 1G, Supplementary data 4). We then tested other cancer types from TCGA with similar analysis for survival and hazard ratio based on GLUT1/3 combination. We found that Pancreatic Ductal Adenocarcinoma (PAAD), Stomach Adenocarcinoma (STAD), Adrenocortical Carcinoma (ACC), Sarcoma (SARC), Liver Hepatocellular Carcinoma (LIHC), Mesothelioma (MESO) and Head and Neck Squamous Cell Carcinoma (HNSC) have significantly poorer survival and higher hazard risk ratio in high-high patients compared to low-low group (Fig. S5A-G; Fig. S6A-G, Supplementary data 5). Together, these data demonstrate that GLUT1 and GLUT3 are jointly upregulated in LUAD, correlate with advanced stage, and strongly predict poor prognosis, establishing glucose transporter co-expression as a clinically relevant metabolic phenotype.

**Fig. 1.**
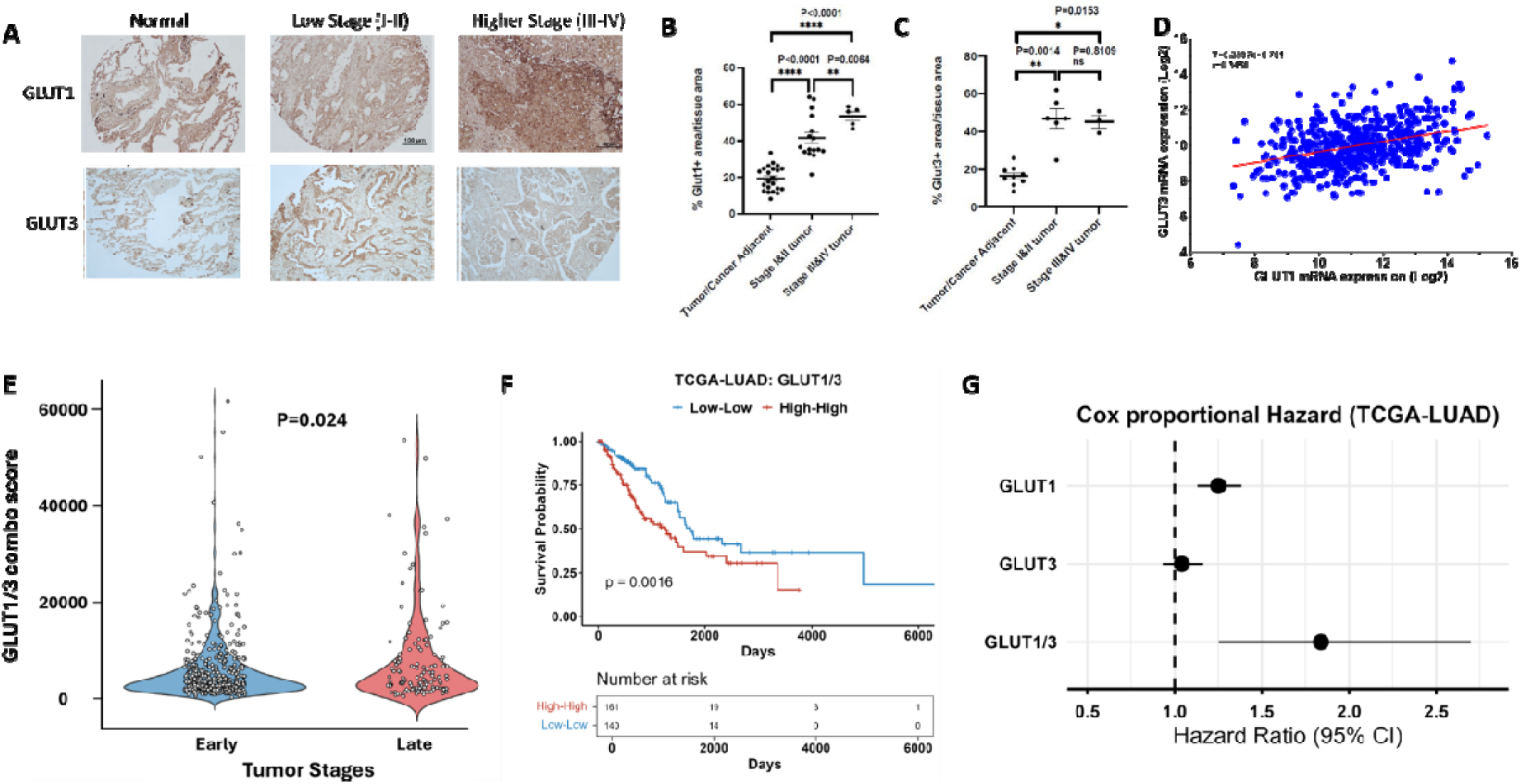
GLUT1 and GLUT3 were prognostic markers for lung adenocarcinoma patients. 1A. Representative IHC analysis of human patient tissue microarray shows comparison between normal lung tissue versus lower stages (Stage I and II) and higher stages (III-IV) for GLUT1 and GLUT3 proteins**. 1B.** Quantification of GLUT1 IHC staining (%GLUT1 area per tissue section) for normal adjacent tissue vs lower stages (I and II) P<0.0001 and lower vs higher stages (III and IV) P=0.0064**. 1C.** Quantification of GLUT3 IHC staining (%GLUT3 area per tissue section) for normal vs lower stages (I and II) P=0.0014 and lower vs higher stages (III and IV) P=0.8109 (Not significant)**. 1D**. Pearson correlation analysis showed positive correlation between GLUT1 and GLUT3 mRNA (R=0.3458; P<0.0001) in expression in TCGA-LUAD cohort. **1E.** GLUT1 and GLUT3 combined score was significantly higher (P=0.024; <0.05) in more aggressive lung cancer patients (Stage III-IC vs Stage I-II) in TCGA-LUAD cohort. **1F**. Kaplan-Meier survival curve of GLUT1/3 shows that high-high patients has poorer survival vs low-low patients (P=0.0049) in TCGA-LUAD cohort. **1G**. Multivariate cox proportional hazard model suggested poorer survival in combined group (high-high vs low-low; HR = 1.72, 95% CI: 1.18–2.51) compared to both individual GLUT1 (high vs low; HR = 1.26, 95% CI: 1.14–1.40) and GLUT3 (high vs low; HR = 1.04, 95% CI: 0.93–1.16) in TCGA-LUAD cohort.

### 3-2. GLUT1 and GLUT3 overexpression combinatorically alter several pathways in LUAD

We observed that combined GLUT1 and GLUT3 score had prognostic power due to positive correlation in LUAD patients (Fig. 1). So, we next tested how the two genes would alter the gene expression as well as biological pathways in LUAD patients. We first started by performing differential gene expression (DEG) analysis and then identified genes that were significantly upregulated (*n*=1645) and downregulated (*n*=1955) for a total of 3600 DEGs (Fig. 2A, Supplementary data 6). We then generated volcano plot illustrates a broad shift toward upregulated genes, with several top hits associated with cell invasion, immune modulation, and tumor metabolism (Fig. 2B, Supplementary data 6). We then performed Gene set enrichment analysis (GSEA) and then further did Hallmark enrichment pathway analysis to identify key pathways altered by the combined score. We found that high-high tumors were enriched for pathways linked tumor aggressiveness and metabolic adaptation, including EMT, E2F targets, TNFα/NF-κB signaling, MYC targets, G2M checkpoint, mTORC1 signaling, inflammatory response, hypoxia, and apoptosis, as well as IFNγ and complement activation (Fig. 2C, Supplementary data 6). These results suggest that high GLUT1/3 expression marks a biologically distinct subtype with enhanced proliferative, inflammatory, and metabolic stress–response programs. We then questioned how the addition of GLUT3 was actually contributing to GLUT1 activity in the combined score compared to GLUT1 alone and so performed variance (ΔR²) across key oncogenic pathways. Adding GLUT3 significantly improved prediction of apoptosis, EMT, hypoxia, and drug-resistance pathway activity, whereas glycolysis showed minimal additional variance (Fig. 2D, Supplementary data 6). This indicates that GLUT3 provides non-redundant biological information distinct from GLUT1. Effect size analysis further confirmed strong pathway-level divergence between groups, with the largest differences observed in hypoxia, glycolysis, and drug-resistance signatures (Fig. 2E, Supplementary data 6). Consistent with these observations, ssGSEA revealed significantly higher activity of EMT, hypoxia, glycolysis, apoptosis, and drug-resistance pathways in High-High tumors (all *P* < 0.0001) (Fig. 2F). We then performed survival analysis based on GLUT1/3 high-high vs low-low groups in GEO dataset GSE68465 to further validate the findings from TCGA-LUAD cohorts (Fig. S7A-B, Supplementary data 7). Heatmaps were also generated for top50 DEGs in TCGA-LUAD data as well as GSE68465 (Fig. 8A-B respectively, Supplementary data 7). Taken together, these results demonstrate that GLUT1/3 co-overexpression identifies a transcriptionally aggressive LUAD subtype characterized by metabolic activation, enhanced stress survival, inflammatory signaling, and resistance-associated gene programs.

### 3-3. DRB18 is a more stable and potent anticancer compound than WZB117

Since, GLUT1 and GLUT3 both contributed to survival and subsequent downstream alteration of different pathways, we further performed experimental analysis with small molecules that targeted both these proteins, our pan class-I GLUT inbhitors. Previously, we reported two glucose transporter inhibitor compounds (Fig. 3A) WZB117 [28–30] and DRB18 [35–38], which can target multiple cancers and have demonstrated anticancer potency in pre-clinical studies. To investigate the stability of these compounds, WZB117 and DRB18 were pre-incubated in serum-containing cell culture media for 0-72 hours and then A549 cells were treated with the compound’s solutions for 24 hours. While DRB18 retained its antiproliferative potency for the full duration of 72 hours, WZB117 showed decreased potency at approximately 12 hours, and completely lost its anticancer activity after 48 hours (Fig. 3B-C). Furthermore, DRB18 had a much higher proliferation-inhibitory activity at 10 mM against 9 major cancer types compared to WZB117 (Fig. 3D). Since DRB18 was a more potent multi-GLUT-targeting compound, we focused on DRB18 in this study. DRB18 reduced cell viability in A549 cells in a time-dependent manner (Fig. 3E). Furthermore, we found that DRB18 reduced glucose uptake in many different cell lines in a dose-dependent manner (Fig. 3F). Further, we tested the acceptability of DRB18 as therapeutic agent by using swissadme portal (http://www.swissadme.ch/index.php). We found that DRB18 had a bioavailability score of 0.55 and drug likeliness based on Lipinski score and high GI absorption. These suggest DRB18 is a possible drug candidate (Supplementary Table 5).

**Fig. 2.**
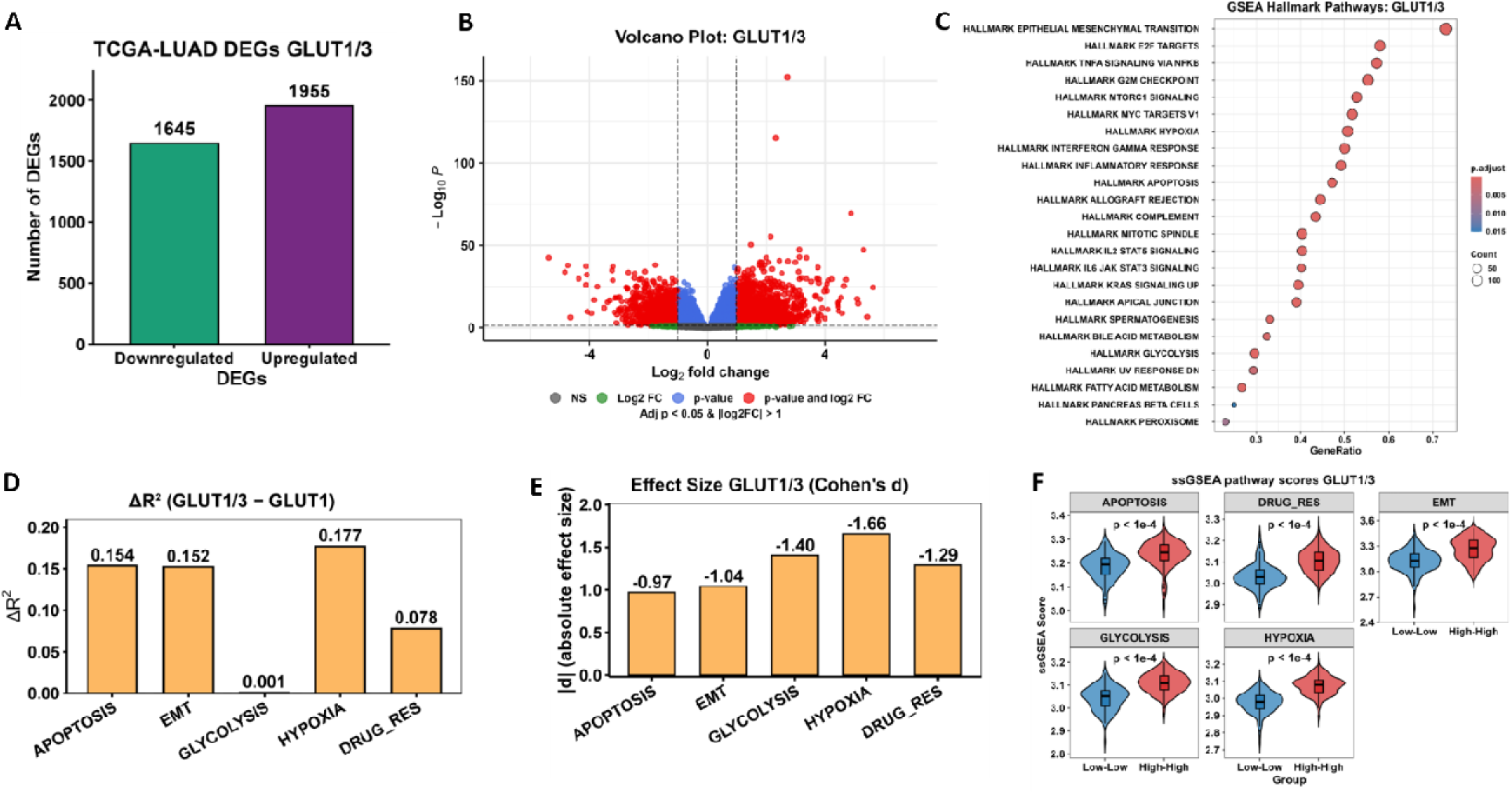
**GLUT1 and GLUT3 combinatorically alter transcriptomics and major biological pathways in LUAD. 2A**. Bar plot summarizing the number of differentially expressed genes (DEGs) between GLUT1/3 High-High and Low-Low LUAD tumors. A total of 1645 genes were downregulated, while 1955 genes were upregulated. **2B.** Volcano plot showing the distribution of DEGs, with color-coding based on significance and magnitude of fold change. Significantly altered genes (adjusted P < 0.05 and |log FC| > 1) are highlighted in red. The top five most significantly upregulated and downregulated genes are labeled. **2C.** Hallmark GSEA comparing GLUT1/3 High-High vs Low-Low tumors. Multiple oncogenic and stress-adaptive pathways were significantly enriched in the High-High group (FDR < 0.05), including EMT, E2F targets TNFα/NF-κB signaling, MYC targets, G2M checkpoint, mTORC1 signaling, hypoxia, apoptosis, and IFNγ pathway. Bubble size represents gene-set size and color denotes adjusted *p*-value. **2D.** Change in explained variance (ΔR²) when adding GLUT3 expression to GLUT1 for predicting pathway scores. GLUT3 significantly increased variance explained for apoptosis (ΔR² = 0.154), EMT (ΔR² = 0.152), hypoxia (ΔR² = 0.177), and drug-resistance pathways (ΔR² = 0.078) (all *P* < 0.05), while contributing minimally to glycolysis. **2E.** Effect sizes (Cohen’s *d*) of ssGSEA pathway score differences between GLUT1/3 High-High and Low-Low tumors. Large effects were observed for hypoxia (|d| = 1.66), followed by drug-resistance (1.29), glycolysis (1.40), EMT (1.04), and apoptosis (0.97), indicating strong biological divergence. **2F.** Violin plots of ssGSEA pathway activity scores in High-High versus Low-Low tumors. GLUT1/3 High-High tumors exhibited significantly elevated pathway activity for apoptosis, EMT, glycolysis, hypoxia, and drug-resistance signatures (all *P* < 0.0001), supporting coordinated activation of metabolic and aggressive programs.

**Fig. 3.**
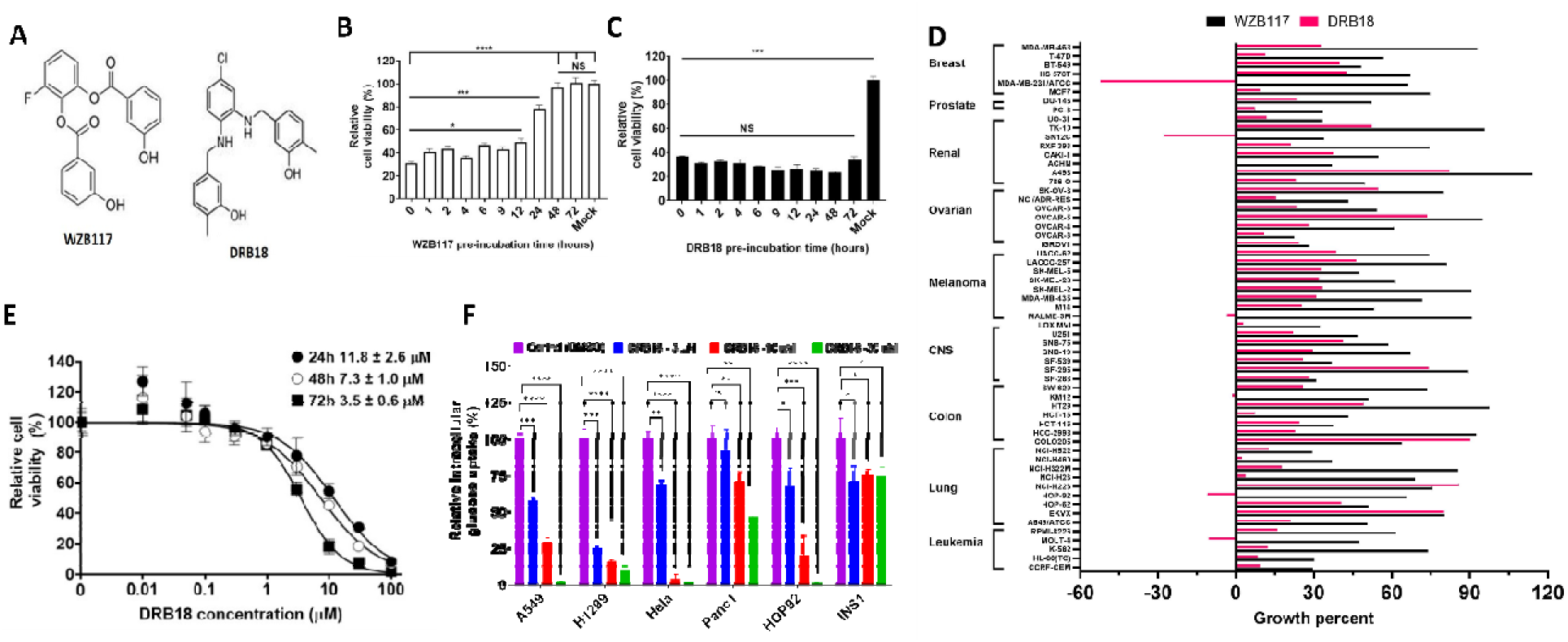
**DRB18 was a more stable and more potent anticancer compound than WZB117. 3A**. Structures of WZB117 and DRB18. **3B**. Potency of WZB117 against NSCLC A549 cells decreased after 12 hours of pre-incubation in serum-containing media. **3C**. Potency of DRB18 against NSCLC A549 cells did not change after 72 hours of pre-incubation in serum-containing media. **3D**. Comparison of potencies between DRB18 and WZB117 in NCI60 cancer cell line panel shows that DRB18 was a more potent anticancer compound compared to WZB117. **3E**. DRB18 reduced the viability in A549 NSCLC cells after 24, 48 and 72 hours of treatment in dose-dependent manner. **3F**. DRB18 reduced intracellular glucose uptake in A549, H1299, Hela, Panc1, HOP92 and INS1 cancer cells in a dose-dependent manner.

### 3-4. DRB18 potency in cancer cells is affected by the availability of extracellular nutrients

To investigate if the potency of DRB18 can be altered by the presence of extracellular nutrients, which directly or indirectly affect central carbon metabolism, we performed different *in vitro* assays in A549, Panc1, and Hela cells, with varying concentrations of glucose, glutamine, and ATP. We firstly found that intracellular ATP levels were dependent on extracellular glucose and extracellular ATP (eATP). Furthermore, DRB18 reduced intracellular ATP (iATP) in dose-dependent, and its activity also depended on extracellular glucose and eATP (Fig. 4A-C). Next, when we treated A549 cells with varying concentrations of glucose in the presence and absence of 10 µM DRB18, and we found that the potency of DRB18 inhibition increased in the presence of higher glucose concentration and further quantification of inhibitory differences showed that DRB18 cytotoxicity towards A549 cells depended directly on extracellular glucose (Fig. 4D-E). Next, we repeated the same experiment in different concentrations of extracellular glutamine and found that extracellular glutamine did not have any significant effect on DRB18 potency but 0 mM glutamine drastically reduced A549 cell viability (Fig. 4F). Similar experiments were performed using extracellular glucose and glutamine in Panc1 cells. We found that extracellular glucose, but not glutamine, had an impact on DRB18 cytotoxicity towards Panc1 cells (Fig. 4G-I). These analyses suggest that DRB18 mediated inhibition of cell proliferation via inhibition of glucose transport, is dependent on eATP and extracellular glucose, and relies to a lesser extent on extracellular glutamine.

**Fig. 4.**
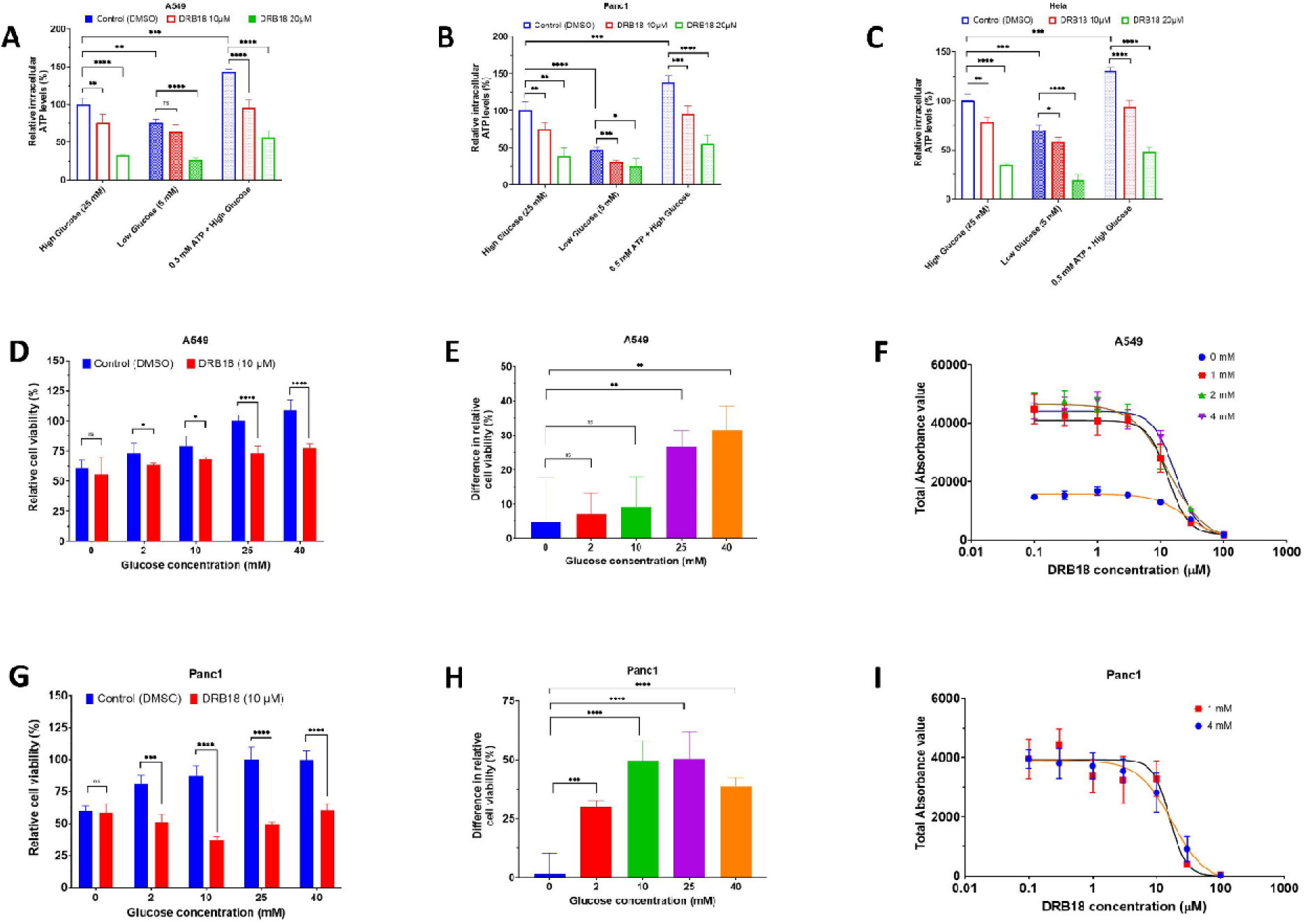
DRB18 potency changed with alterations in extracellular nutrients *in vitro* in A549 and Panc1 cancer cells. 4A. Relative intracellular ATP (iATP) changes with increasing concentrations of DRB18 (0, 10 and 20 µM) under conditions of high glucose (25 mM), low glucose (5 mM) and 0.5 mM extracellular ATP (eATP) in A549 cells. **4B.** Relative iATP changes with increasing concentrations of DRB18 (0, 10 and 20 µM) under conditions of high glucose (25 mM), low glucose (5 mM) and 0.5 mM eATP in Panc1 cells. **4C.** Relative intracellular iATP changes with increasing concentrations of DRB18 (0, 10 and 20 µM) under conditions of high glucose (25 mM), low glucose (5 mM) and 0.5 mM eATP conditions in Hela cancer cells. **4D**. Relative cell viability in A549 NSCLC cells after 10 µM DRB18 treatment with varying concentrations of glucose (0, 2, 10, 25 and 40 mM). **4E**. DRB18 exhibited significantly more differences in viability in A549 cells at 10-, 25- and 40-mM glucose concentrations compared to 0- and 2-mM extracellular glucose. **4F**. DRB18 inhibited total absorbance (cell viability) in A549 cells in a dose-dependent manner with varying concentrations of extracellular glutamine (0, 1, 2 and 4 mM). **4G**. Relative cell viability in Panc1 pancreatic cancer cells after 10 µM DRB18 treatment with varying concentrations of glucose (0, 2, 10, 25 and 40 mM). **4H**. DRB18 exhibited significantly more differences in viability in Panc1 cells at 10-, 25- and 40-mM glucose concentrations compared to 0- and 2-mM extracellular glucose. **4I**. DRB18 inhibited total absorbance (cell viability) in Panc1 cells in a dose-dependent manner with varying concentrations of extracellular glutamine (1, 4 mM).

### 3-5. DRB18 exhibits synergistic effects with anticancer compounds *in vitro*

To investigate the clinical applicability of DRB18, we tested synergistic effects of the compound with other anticancer drugs, which were either in clinical use or used in pre-clinical models. We found that glucose and glutamine can synergistically reduce cell viability in A549 and Panc1 cells (Fig. S9A and S9B). Based on these results, the glutamine transporter inhibitor V9302 and glutaminase I inhibitor CB839 were used in both the cell lines. We first determined the IC_50_ of V9302 and CB839 in both cell lines (Fig. S9C-F). We found that the combination of DRB18 and V9302 showed more inhibitory activity towards A549 and Panc1 cells in comparison to no treatment or single treatment of both compounds (Fig. 5A-D). Similarly, dual treatment of DRB18 and CB839 showed significantly more cytotoxicity in A549 and Panc1 cells (Fig. 5B and 5E). Sunitinib is an FDA approved receptor tyrosine kinase inhibitor in different cancer types. We found that DRB18 exhibited additive effect with sunitinib in both A549 and Panc1 cells (Fig. 5C and 5F). Brigatnib is an FDA approved drug for metastatic NSCLC. We determined the IC_50_ of brigatinib in A549 cells (Fig. S9G). We further found that DRB18 showed no additive effect with brigatinib, but the combined treatment of DRB18 and brigatinib was able to reduce cell viability in A549 cells significantly more than control or either compound working alone (Fig. 5G). Trametinib is an FDA approved drug for treatment of melanoma and NSCLC and is a MEK inhibitor. We determined the IC_50_ of trametinib in A549 cells (Fig. S9H) and found that DRB18 and 15 µM of trametinib exhibited additive effects in A549 cells, although no such effect was observed between DRB18 and 10 µM trametinib (Fig. 5H).

**Fig. 5.**
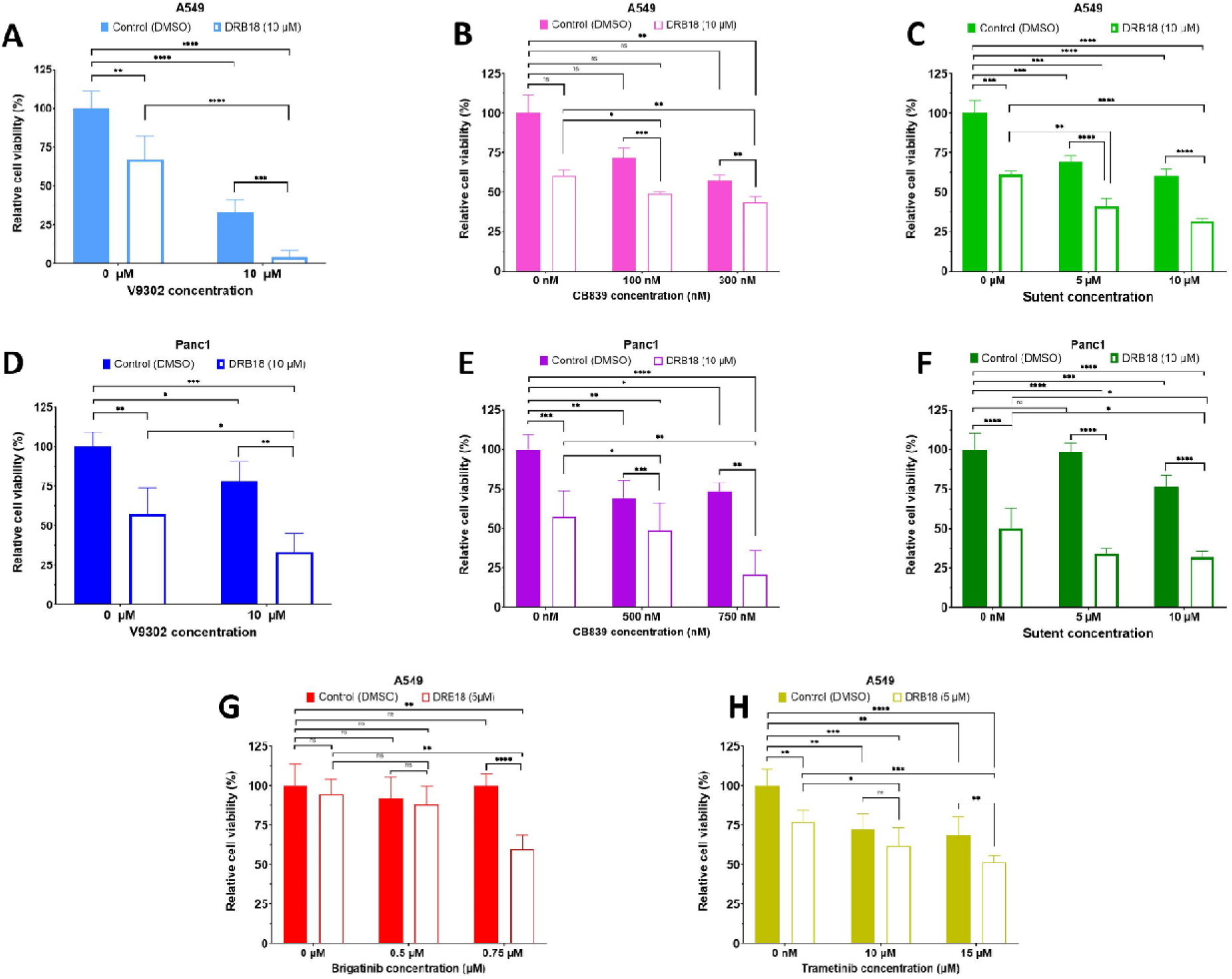
**DRB18 exhibited additive anti-proliferative effects against different cancer cell lines with several clinical and pre-clinical anticancer agents *in vitro.*** 10 µM of DRB18 combined with or without other drugs to treat A549 cells for 48 hours. The inhibition of cell proliferation was compared to single compound treatment to identify anti-proliferation synergism. **5A.** DRB18 combined with 10 µM V9302 showed more inhibition against A549 cells. **5B**. DRB18 combined with 100 nM or 300 nM of CB839 showed more cytotoxicity against A549. **5C**. DRB18 combined with 5 µM or 10 µM of sunitinib (Sutent) showed more inhibition against A549 cells. **5D**. DRB18 combined with 10 µM V9302 showed more inhibition against Panc1 cells. **5E**. DRB18 combined with 100 nM or 300 nM of CB839 showed more inhibition against Panc1 cells. **5F**. DRB18 combined with 5 µM or 10 µM of sunitinib showed more inhibition against Panc1 cells. **5G**. 5 µM of DRB18 combined with 0.75 µM of brigatinib showed more cytotoxicity against A549 cells. **5H**. 5 µM of DRB18 combined with 15 µM of trametinib showed more inhibition against A549 cells.

### 3-6. DRB18 exhibits synergistic effects with Paclitaxel *in vitro*

Paclitaxel is an FDA approved drug for metastatic NSCLC and is known to target tubulin polymerization in lung cancer cells [40–42]. We tested the potential synergism of DRB18 with paclitaxel in A549 and Panc1 cells. We found that 10 µM DRB18 and 10 µM paclitaxel showed synergism by inhibiting cell growth in A549 cells after 48 hours of treatment with varying dosages of paclitaxel (Fig. 6A-C). We computed synergism by using synergyfinder (synergyfinder.org) and used an HS model to determine the synergism score of 17.56 (P ≤ 0.001). We tested administration of 10 µM DRB18 and 10 µM Paclitaxel in Panc1 cells and found that dual treatment reduced cell growth significantly after 48 hours, compared to either single compound or control (Fig. 6D). Since DRB18 reduces glucose transport and leads to glucose deprivation in cancer cells, we wanted to determine whether low glucose conditions could exhibit synergism with paclitaxel in A549 and Panc1 cells. We found that dual treatment of 1 µM paclitaxel and low glucose (2 mM) conditions inhibited cell growth in A549 cells more significantly than either condition alone (Fig. 6E). This effect was not observed at 0.3 µM paclitaxel concentration. However, in Panc1 cells, both 0.3 and 1 µM paclitaxel in low glucose (2 mM) conditions were able to reduce cell growth significantly more than in either condition alone (Fig. 6F).

**Fig. 6.**
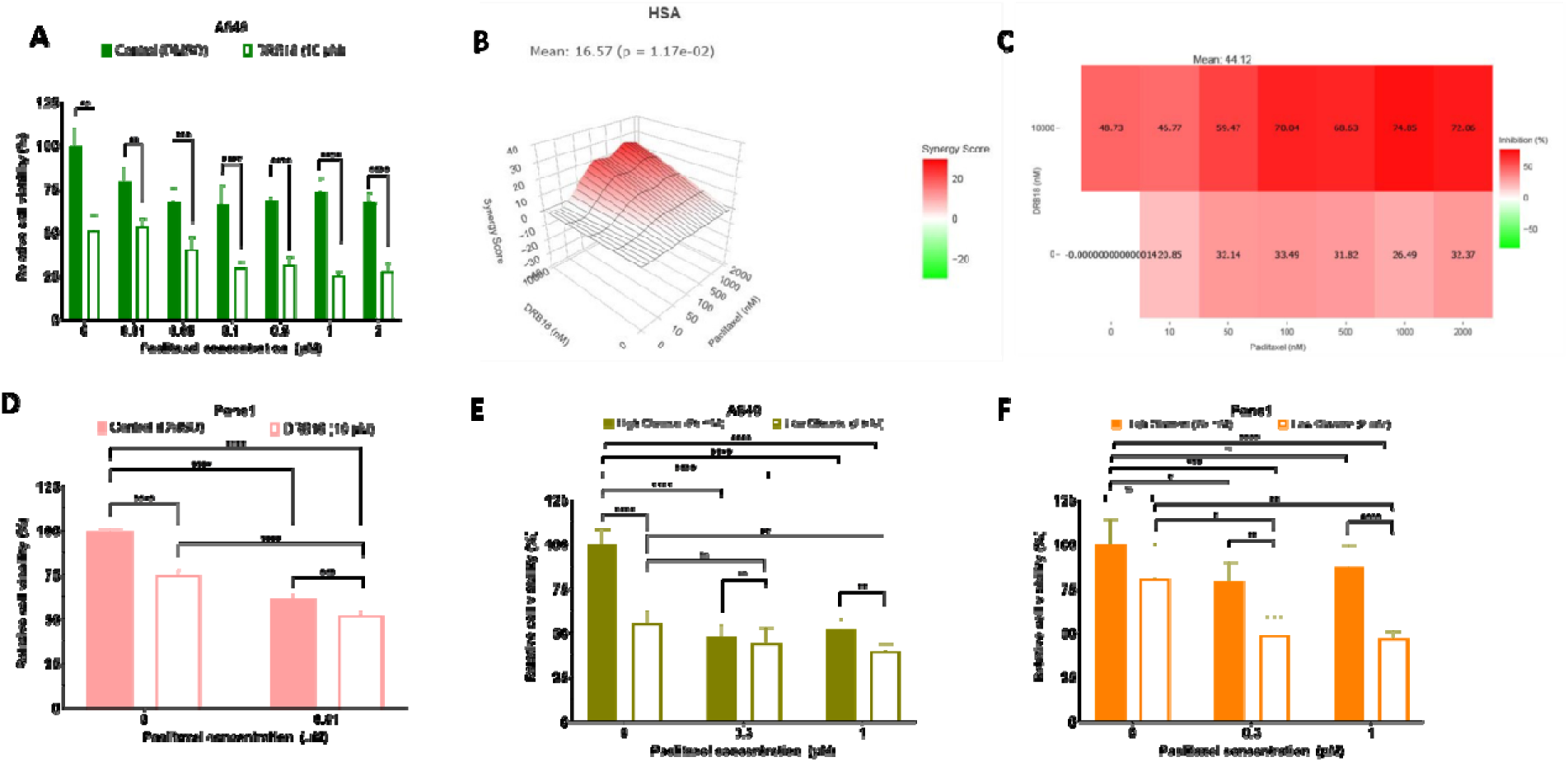
DRB18 exhibited anti-proliferative synergism with clinical agent paclitaxel *in vitro.* 6A. DRB18 exhibits synergism with different dosages of anticancer compound paclitaxel. **6B.** HSA model obtained synergism score of 16.57 (P-value of 0.017). **6C**. Synergistic analysis showed a synergy mean value of 44.12 indicating high synergy scores between the DRB18 and paclitaxel. **6D**. DRB18 combined with 100 nM of paclitaxel significantly higher anti-proliferative effect against Panc1 cells. **6E**. Low glucose (2 mM) treatment combined with 1 µM paclitaxel showed a higher anti-proliferative effect against A549 cells. **6F**. Low glucose (2 mM) treatment combined with 0.3 and 1 µM paclitaxel showed a significantly higher anti-proliferative effect against Panc1 cells.

### 3-7. DRB18 exhibits synergistic effects with paclitaxel *in vivo*

We next wanted to test if the combination of DRB18 and paclitaxel was synergistic *in vivo*. Nude mice were implanted with subcutaneous injections of A549 cells and then were treated with either vehicle, 8 mg/kg DRB18, 8 mg/kg paclitaxel, or combined DRB18+paclitaxel for 5 weeks. We found that dual treatment of DRB18 and paclitaxel significantly reduced the tumor volume by approximately 79%, 73%, and 50% in comparison to vehicle, DRB18 alone, and paclitaxel alone, respectively (Fig. 7A-B). Comparing the final tumor weights, they were reduced by approximately 87%, 85%, and 71% in comparison to vehicles, DRB18 alone, and paclitaxel alone, respectively (Fig. 7C). We also observed that there was no significant reduction in the average body weight of the mice in the four groups as well as no differences in food consumption over the 5-week span from the start of the treatment (Fig. S10A-B).

**Fig. 7.**
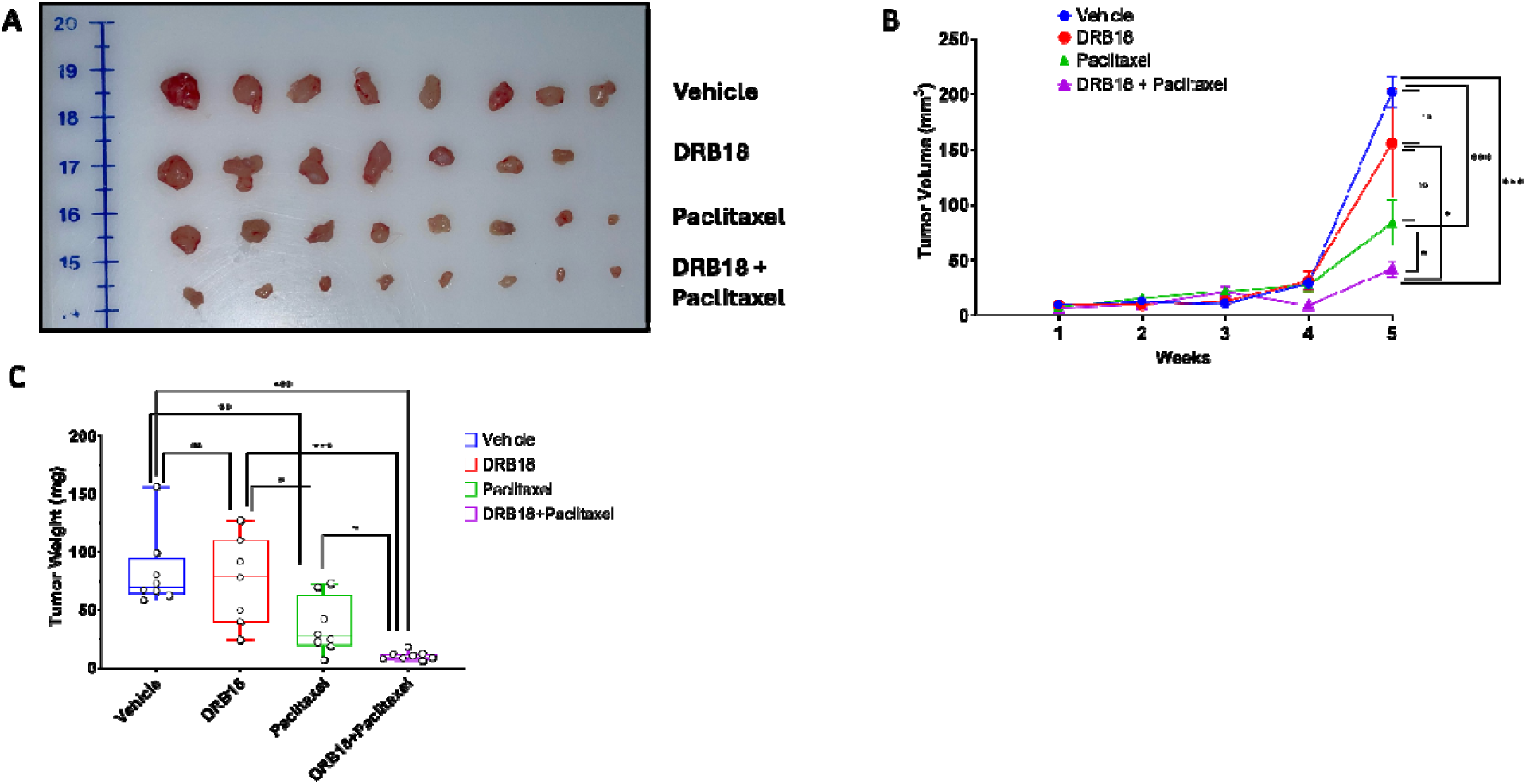
**DRB18 exhibited synergism of anti-tumor growth with NSCLC anticancer drug paclitaxel in vivo. 7A**. Schematic images of tumors obtained after mice were euthanized after 5 weeks of treatment. **7B**. Combined treatment of DRB18 and paclitaxel reduced tumor volume compared to either DRB18 or paclitaxel alone or vehicle treatment. **7C**. Combined treatment of DRB18 and paclitaxel reduced tumor weight compared to either DRB18 or paclitaxel alone or vehicle treatment.

## Discussion

### Most cancers upregulate class I GLUTs, and DRB18 is a pan-class I GLUT inhibitor

GLUTs play critical roles in cellular homeostasis by ensuring sufficient glucose supply as the main carbon and energy source for mammalian cells [14–16]. Class I GLUTs are frequently upregulated across cancer types and have been linked to proliferation, drug resistance and metastasis [19, 20, 43]. We previously showed that the pan–class I GLUT inhibitor DRB18 reduces GLUT-mediated glucose transport, energy metabolism and cell viability in human lung cancer models in vitro and in vivo [38]. In the present study, we extended these observations by systematically analyzing the expression patterns and clinical impact of GLUT1 and GLUT3 in LUAD patients, and by defining the transcriptional programs associated with combined high GLUT1/3 expression.

Immunohistochemistry of human LUAD tissue microarrays showed that both GLUT1 and GLUT3 protein levels were significantly increased in LUAD samples compared with adjacent normal lung tissue (Fig. 1A–C). Notably, GLUT1 expression further increased from early- to late-stage disease, whereas GLUT3 levels were change significantly with further stage progression (Fig. 1B–C). In TCGA-LUAD, a combined GLUT1/3 expression score was higher in stage III/IV tumors, whereas only GLUT1 mRNA alone showed a clear gradient with stage and nodal status (Fig. 1E; Fig. S1E–H). Subsequently, TCGA-LUAD patients were divided into GLUT1 and GLUT3 (High-High group and low-low group). Survival analysis revealed that GLUT1/3 high-high group had poorer overall survival compared with Low-Low patients (Fig. 1F) and Cox hazard analysis revealed that GLUT1/3 group showed the largest hazard ratio in comparison to either individual GLUT1 or GLUT3 (Fig. 1G). These analyses were found in other cancer types as well suggesting that GLUT1/3 combination had prognostic value in many but not all cancer types (Fig. S4-5). In line with these transcript-level findings, GLUT1 and GLUT3 proteins were positively correlated in CPTAC LUAD specimens and across lung cancer cell lines in CCLE, and this positive correlation extended to multiple other tumor types (Fig. S2A–G). Collectively, these data indicate that while GLUT1 is the dominant individual prognostic transporter in LUAD, co-upregulation of GLUT1 and GLUT3 identifies a particularly aggressive metabolic phenotype.

To understand the biological programs associated with GLUT1/3 co-overexpression, we compared the transcriptomes of GLUT1/3 High-High versus Low-Low LUAD tumors. This analysis revealed >3,500 DEGs, indicating extensive transcriptional reprogramming associated with combined high GLUT1 and GLUT3 (Fig. 2A–B). Hallmark GSEA showed strong enrichment of EMT, E2F targets, hypoxia, TNFα/NF-κB signaling, G2M checkpoint and apoptosis pathways in the High-High group (Fig. 2C), all of which are linked to aggressive tumor behavior, treatment resistance and metabolic stress adaptation. Importantly, variance decomposition analyses demonstrated that adding GLUT3 to GLUT1 significantly improved the prediction of EMT, hypoxia, apoptosis and drug-resistance pathway activity, whereas glycolysis depended predominantly on GLUT1 alone (Fig. 2D). Consistently, ssGSEA effect sizes were largest for hypoxia, glycolysis and drug-resistance signatures, and pathway scores were markedly higher in High-High tumors than in Low-Low tumors (Fig. 2E–F). Further validation was obtained using GSE68465 as LUAD microarray for survival analysis (Fig. S7-S8). These findings suggest that GLUT3 does not simply mirror GLUT1 expression but rather amplifies specific stress-adaptive and survival pathways in GLUT1-high tumors, which may explain the particularly poor prognosis of the GLUT1/3 High-High subgroup.

Although inhibiting glucose transport and glucose-mediated metabolism, in theory, should be effective in suppressing tumor growth, most cancer cells express more than one GLUT [44,45], and different GLUT isoforms are known to play distinct roles in tumor growth and metastasis. We recently reported that DRB18 is a pan-class I GLUT inhibitor that acts on GLUT1–4 with variable inhibitory potency [38]. In addition, tumors generated from A549 cells lacking GLUT1 (A549 GLUT1-knockout cells) grew as fast as tumors generated from parental A549 cells [38]. These results suggest that other class I GLUTs (GLUT2–4) can compensate for the loss of GLUT1 in A549-GLUT1KO tumors. In contrast, DRB18 inhibited A549 tumor growth due to its pan–class I GLUT-targeting activity [38].

### Cancers use alternative pathways to bypass drug treatment, and combinational therapy with DRB18 can overcome this problem

Cancers have their built-in flexibility in bypassing enzyme / pathway inhibitions and generating resistance to a treatment with mono anticancer therapy. A combinatorial therapy with two anticancer drugs, including a GLUT inhibitor such as DRB18, will be much more advantageous over single drug therapies when the cancer growth relies on glucose transport and on another cell-growth signaling pathways that are unrelated to glucose transport / glucose metabolism.

Personalized monotherapy using chemo or targeted drugs are still prevalent in the clinic. However, the major concern associated with single drug therapy is toxicity due to higher drug dosages and induced drug resistance, resulting in tumor relapses after initial remission. Cancer cells are dependent on multiple mechanisms for survival. It becomes necessary, therefore, to target multiple mechanisms simultaneously for better therapeutic outcomes. Thus, identifying and using combo therapy in a synergistic combination is imperative to reduce the burden of high drug dosages as well as undesired side-effects of each drug in a drug pair.

### Anticancer synergism between DRB18 and paclitaxel

The main criterium for synergism between two drugs is that the anticancer of two drugs working together can achieve a superior efficacy to the total sum of the individual anticancer efficacies of the drug pair working alone. Synergy is relatively rare to find between two anticancer agents and is difficult to predict *in vitro* and even more so *in vivo*.

In this study, we report the additive or synergistic effects of DRB18 along with other FDA approved or clinically tested anticancer drugs. Investigation of DRB18 properties revealed (http://www.swissadme.ch, 04 January 2024) that DRB18 has a positive Lepinski score, good gastrointestinal absorption, and a high bioavailability score of 0.55, further justifying its potential as a drug candidate (Supplementary Table 5). One of the major objectives of this study was to investigate the synergistic mechanisms of DRB18 with other clinical or preclinical anticancer agents. Among anticancer drugs and compounds tested, we found paclitaxel to be the most suitable candidate for forming a synergistic pair with DRB18. Paclitaxel is the gold standard chemotherapy drug for advanced NSCLC and for patients who are sensitive to immunotherapy. Paclitaxel has several properties to make it suitable as a drug candidate, but sometimes monotherapy fails due to drug induced toxicity such as neurotoxicity [46–48].

The observation that GLUT1/3 High-High tumors are enriched for hypoxia, glycolysis, EMT and drug-resistance programs (Fig. 2) provides a mechanistic rationale for targeting class I GLUTs with DRB18 in combination with cytotoxic agents such as paclitaxel. By simultaneously disrupting glucose-uptake–driven metabolic plasticity and microtubule dynamics, the DRB18–paclitaxel combination may be particularly effective against this metabolically rewired, treatment-resistant LUAD subtype. From a clinical perspective, such a combination could allow dose reduction of paclitaxel while maintaining or enhancing anticancer efficacy.

During tumor treatment, no adverse effects were observed, including no differences in food intake and body weight (Fig. S10A-B) in the DRB18+paclitaxel group, compared with the DRB18 group or paclitaxel group. This might be due to dosage and/or administration intervals of the drugs we used. In our previous tumor studies, 10 mg/kg was used for DRB18 [38], whereas the generally used dosage for paclitaxel is 10 mg/kg. In the pilot and full-scale studies of this investigation, 8 mg/kg for both DRB18 and paclitaxel was used and found to be effective in reducing tumor growth (Fig. 7), without adverse effects on body weight and food intake. Additional fine-tuning adjustments in dosages and drug administration frequency may further improve the treatment results.

## Conclusions

Taken together, our data demonstrated that human lung cancers overexpress GLUT1 and GLUT3 where they work combinatorically to reduce survival by altering many biological pathways including drug resistance, while our pan-class I GLUT inhibitor DRB18 inhibits GLUT1-4 (36). Thus, DRB18 is a wide spectrum inhibitor for class I GLUTs and GLUT-upregulating NSCLC and other cancers. While working alone, DRB18 reduced tumor size of NSCLC A549 cells. The combinatorial therapy of DRB18 and paclitaxel further reduced the size of tumors significantly, showing anticancer synergy. Moreover, these adjustments led to no significant changes in food intake and body weight, compared with mock-treated controls. Thus, DRB18, as a pan-class I GLUT inhibitor, can function alone or in combination with other approved anticancer drugs to treat different cancers, as paclitaxel has been approved for treating eight different cancers, to achieve greater anticancer efficacy. Additionally, more approved anticancer drugs will likely be identified to exhibit synergy with DRB18, further widening the applications of DRB18 or pan class I GLUT inhibitors in cancer therapies. These may significantly increase the use and values of the pan class I GLUT inhibitors in new cancer therapies. Noteworthy, GLUT4 has recently been found implicated in aging and neuronal diseases such as Alzheimer’s and Parkinson’s [49], further expanding DRB18’s utility in treating human diseases beyond cancers.

## List of abbreviations

LUAD: Lung adenocarcinoma
DMSO: Dimethyl sulfoxide
GLUT: Glucose transporter
NSCLC: Non-small cell lung cancer
NS: Not Significant

## Declarations

### Ethics approval and consent to participate

All animal studies described below were conducted in accordance with US government regulation on animal care and Ohio University IACUC approved protocols.

### Consent for publication

Not applicable.

### Availability of data and materials

All the data generated in this study for bioinformatics has been obtained from https://portal.gdc.cancer.gov using TCGAbiolinks package in R. The independent validation cohort GSE68465 (NSCLC microarray dataset with survival data) was downloaded using GEOquery package of R. Data used in correlation analysis has been obtained from LUAD and CCLE databased available in cbioportal (https://www.cbioportal.org/datasets). The data analyzed for the generation of different graphs such as survival plots, volcano plots, DEGs, cox hazard models, GSEA, pathway scores, etc, have been available in supplementary data. The R-code used for performing this study could be obtained from the corresponding author upon generous and sincere request. Data used for generation of animal studies graphs can be provided from corresponding author upon generously and sincerely request.

## Acknowledgments

Not applicable

## Authors’ contributions

P.S. designed and performed most of the experiments. P.S., L.B., R.W and J.S. performed resazurin cell viability experiments. P.S., L.B., H.Z. and Y.L. performed animal experiments. S.B., and D.R. provided the compounds DRB18 and WZB117. S.A. and C.N. conducted IHC analysis of tissue microarray. X.C. provided guidance and shared lab space and equipment for most experiments. P.S. and X.C. wrote the manuscript. The authors read and approved the final manuscript.

Conflict of interest disclosure statement

The authors declare no potential conflicts of interest.

