## supplementary file for "A novel pan class-I glucose transporter inhibitor DRB18 exhibits synergistic effects with paclitaxel *in vitro* and *in vivo* against human non-small cell lung cancer"

**Supplementary Methods, Figure and Tables**

**Bioinformatics analysis**

GLUT1 and GLUT3 protein expression analysis and comparison between grade 2 and grade 3 tumors were performed using UALCAN webserver (<https://ualcan.path.uab.edu>). Both mRNA and protein levels in Cancer cell line encyclopedia (CCLE) dataset (Broad Institute, 2019) in lung cancer cell lines as well as combined cancer cell lines were downloaded from cbioportal. Protein-protein interaction study was performed using genemania (https://genemania.org/) using GLUT1 and GLUT3. Combined interacting partners analysis for GLUT1 and GLUT3 was performed using String database (https://string-db.org/). Gene Ontology (GO) analysis was performed using shinyGOv0.741 (<http://bioinformatics.sdstate.edu/go74/>).

**Figure legends of Supplementary Figures.**

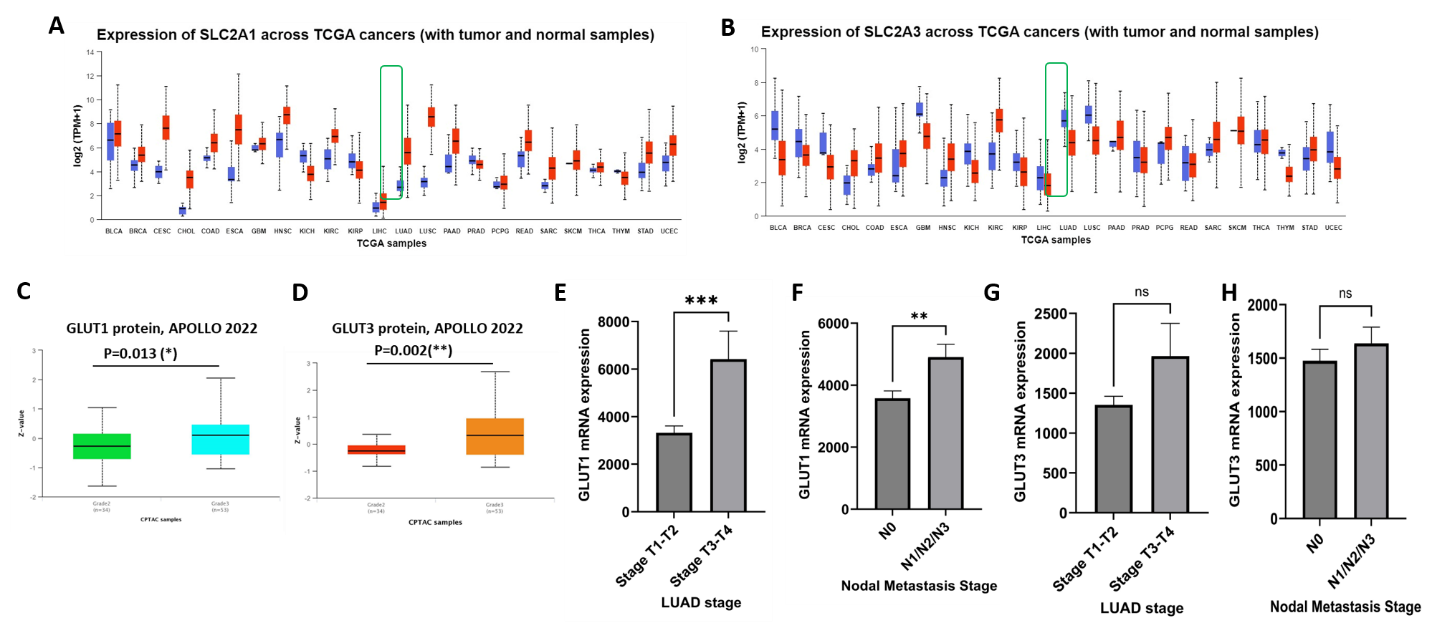

**Figure S1**. **Pan-cancer expression analysis of GLUTs across TCGA cohort**.

**S1A**. *GLUT1* (SLC2A1) expression in TCGA-pan cancer cohort (UALCAN webserver) showed that *GLUT1* mRNA expression was increased in many different cancer types including LUAD compared to normal patients. **S1B**. *GLUT3* (SLC2A3) expression in TCGA-pan cancer cohort (UALCAN webserver) showed that *GLUT3* mRNA expression was increased in many different cancer types including LUAD compared to normal patients. **S1C.** GLUT1 protein expression was significantly increased from grade 2 (n=34) to grade 3 (n=53) tumors in APOLLO LUAD cohort**. S1D.** GLUT3 protein expression was significantly increased from grade 2 (n=34) to grade 3 (n=53) tumors in APOLLO LUAD cohort**. S1E.** *GLUT1* mRNA was increased in patients from lower stages (Stage T1-T2; n=198) to higher stages (Stage T3-T4; n=32) in TCGA-LUAD cohort. **S1F.** *GLUT1* mRNA was increased in patients with no nodal metastasis (N0; n=333) compared to those with nodal metastasis (N1/N2/N3; n=172) in TCGA-LUAD cohort. **S1G.** *GLUT3* mRNA did not increase in patients from lower stages (T1-T2; n=198) to higher stages (T3-T4; n=32) in TCGA-LUAD cohort. **S1H**. *GLUT3* mRNA was increased in patients with no nodal metastasis (N0; n=333) compared to those with nodal metastasis (N1/N2/N3; n=172) in TCGA-LUAD cohort.

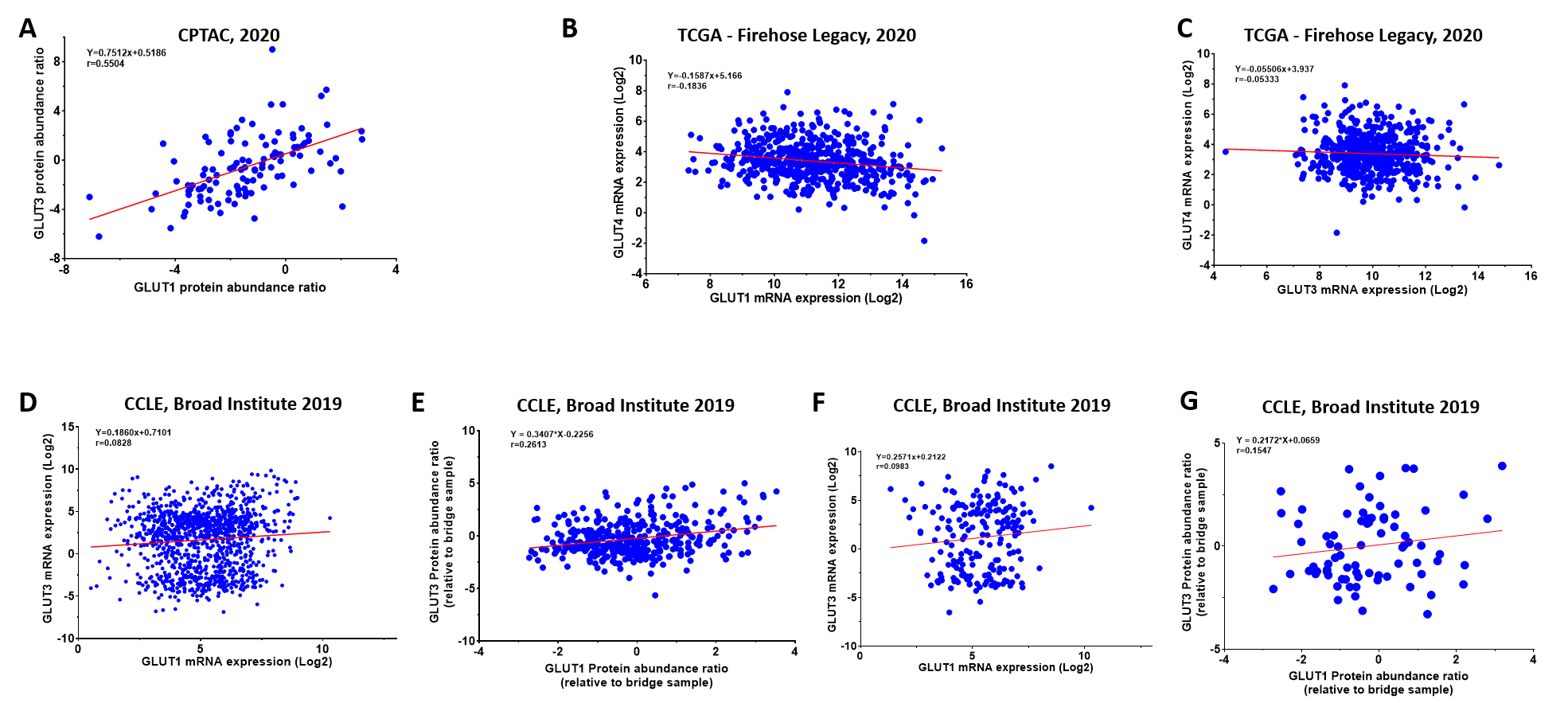

**Figure S2**. **Survival and co-expression analyses of** **GLUT1 and GLUT3 expression.**

**S2A**. GLUT1 and GLUT3 proteins showed positive correlation in expression in CPTAC-LUAD cohort. **S2B**. *GLUT1* and *GLUT4* mRNA showed no significant correlation in expression in LUAD-TCGA cohort. **S2C**. *GLUT3* and *GLUT4* mRNA showed no significant correlation in expression in LUAD-TCGA cohort. **S2D.** *GLUT1* and *GLUT3* mRNA showed positive correlation in expression in lung cancer cell lines (CCLE, Broad Institute dataset, 2019). **S2E**. GLUT1 and GLUT3 proteins showed positive correlation in expression in lung cancer cell lines (CCLE, Broad Institute dataset, 2019). **S2F**. *GLUT1* and *GLUT3* mRNA positively correlated in CCLE dataset (Broad Institute, 2019). **S2G**. GLUT1 and GLUT3 proteins showed positive correlation in expression in CCLE dataset (Broad Institute, 2019).

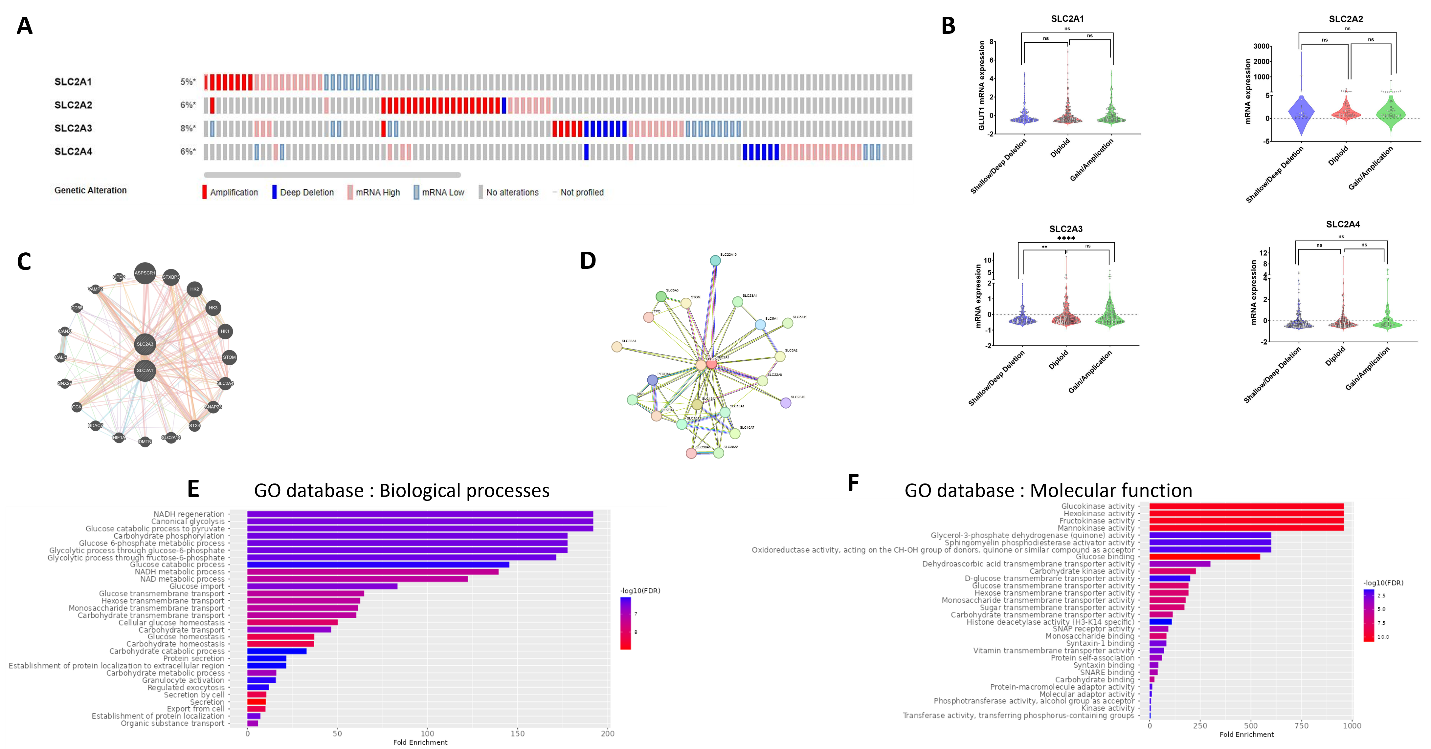

**Figure S3**. **GLUT1 and GLUT3 expression analysis based on copy number alterations, mutations and interacting pathways**

**S3A**. Mutations analysis on GLUT1-4 in TCGA LUAD cohort (oncoprint tool in cbioportal webserver; https://www.cbioportal.org/). **S3B**. Copy number alterations analysis of GLUT1-4 in TCGA LUAD. **S3C**. Genemania interaction analysis of top interacting partners of *GLUT1* and *GLUT3* genes. **S3D**. String database analysis of interacting proteins with GLUT1 and GLUT3. **S3E**. ShinyGO (v0.741) based analysis for gene ontology (biological processes) for top 25 interacting partners for GLUT1 and GLUT3. **S3F**. ShinyGO (v0.741) based analysis for gene ontology (molecular function) for top 25 interacting partners for GLUT1 and GLUT3.

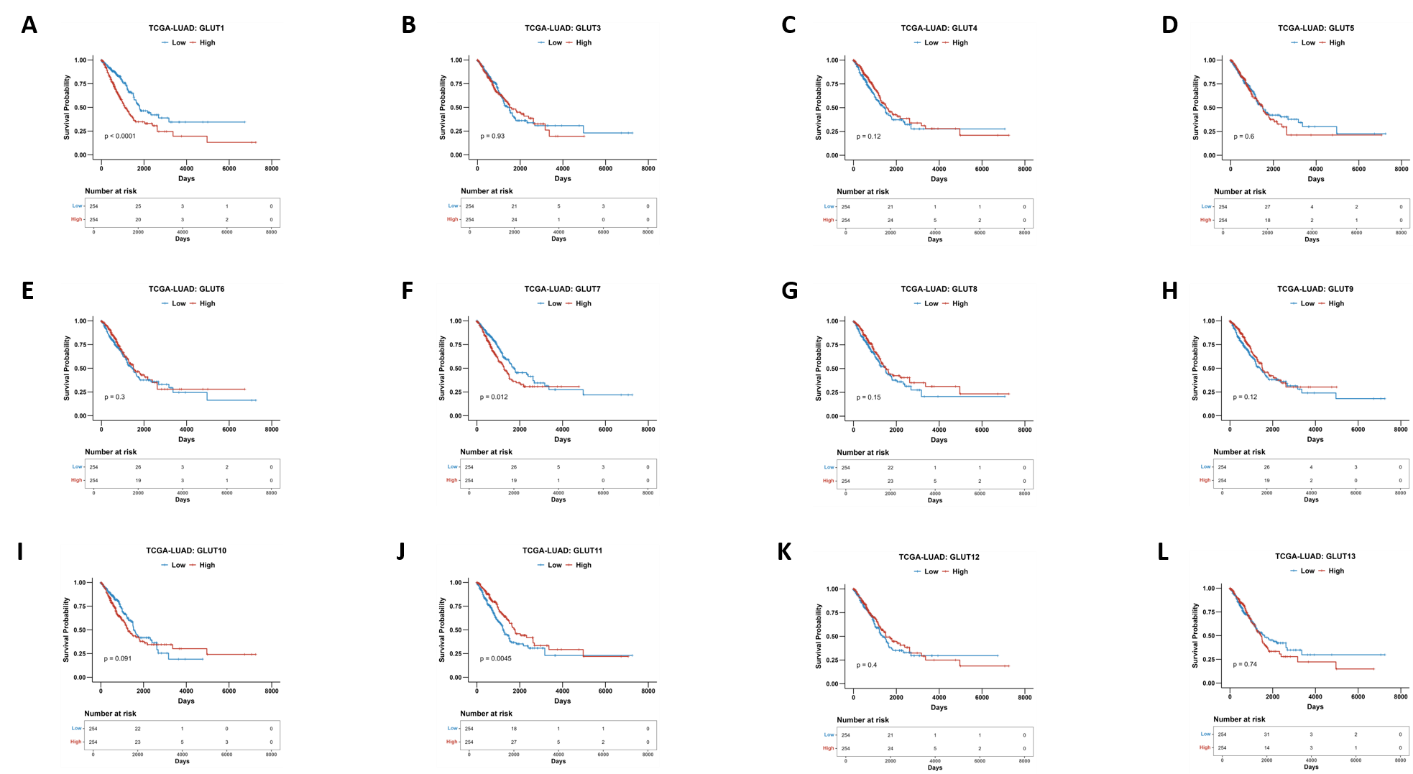

**Figure S4**. **Survival analyses of** **GLUT1-13 in TCGA-LUAD dataset**

**S4A.** Differential overall survival of patients with high or low *GLUT1* mRNA expression. **S4B**. Differential overall survival of patients with high or low *GLUT3* mRNA expression. **S4C**. Differential overall survival of patients with high or low *GLUT4* mRNA expression. **S4D**. Differential overall survival of patients with high or low *GLUT5* mRNA expression. **S4E**. Differential overall survival of patients with high or low *GLUT6* mRNA expression. **S4F**. Differential overall survival of patients with high or low *GLUT7* mRNA expression. **S4G**. Differential overall survival of patients with high or low *GLUT8* mRNA expression. **S4H**. Differential overall survival of patients with high or low *GLUT9* mRNA expression. **S4I**. Differential overall survival of patients with high or low *GLUT10* mRNA expression. **S4J**. Differential overall survival of patients with high or low *GLUT11* mRNA expression. **S4K**. Differential overall survival of patients with high or low *GLUT12* mRNA expression. **S4L**. Differential overall survival of patients with high or low *GLUT1*3 mRNA expression.

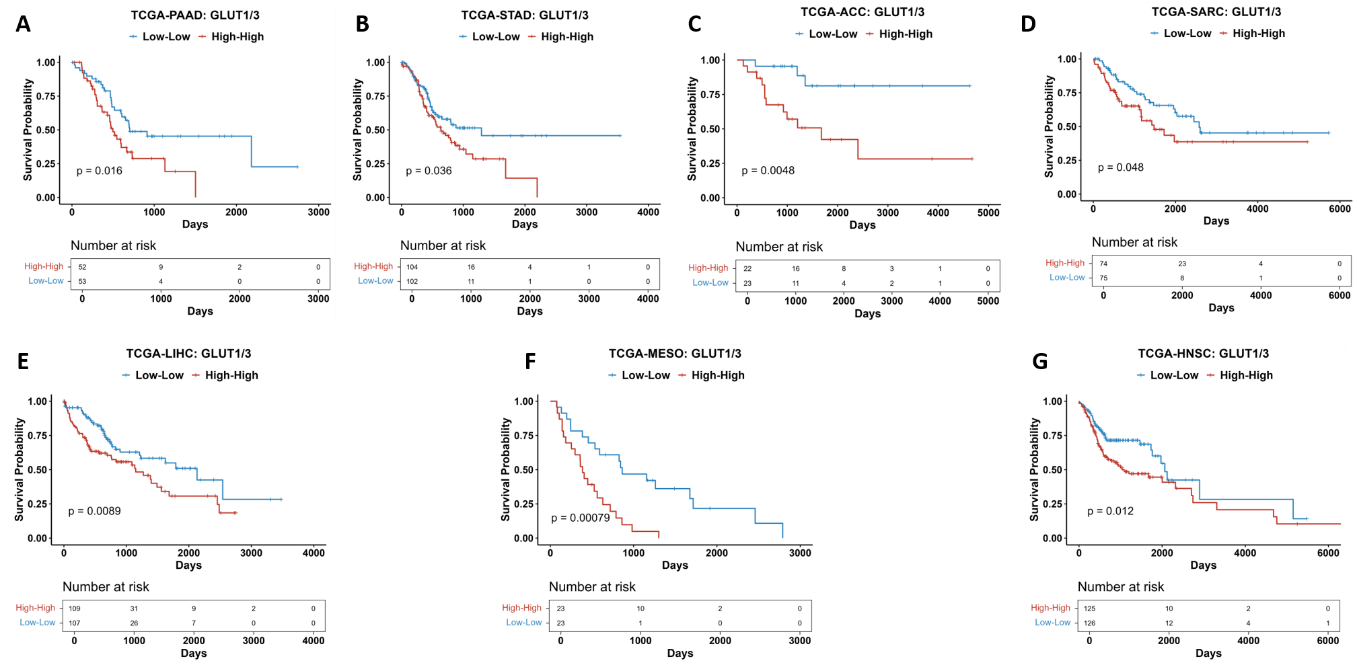

**Figure S5**. **Overall survival analyses of** **patients with GLUT1/3 high-high vs low-low based on mRNA expression in TCGA cancer types. Significant values are shown.**

**S4A**. Overall survival of patients in TCGA-PAAD based on GLUT1/3 expression. **S4B**. Overall survival of patients in TCGA-STAD based on GLUT1/3 expression. **S4C**. Overall survival of patients in TCGA-ACC based on GLUT1/3 expression. **S4D**. Overall survival of patients in TCGA-SARC based on GLUT1/3 expression. **S4E**. Overall survival of patients in TCGA-LIHC based on GLUT1/3 expression. **S2F**. Overall survival of patients in TCGA-MESO based on GLUT1/3 expression. **S4G**. Overall survival of patients in TCGA-HNSC based on GLUT1/3 expression.

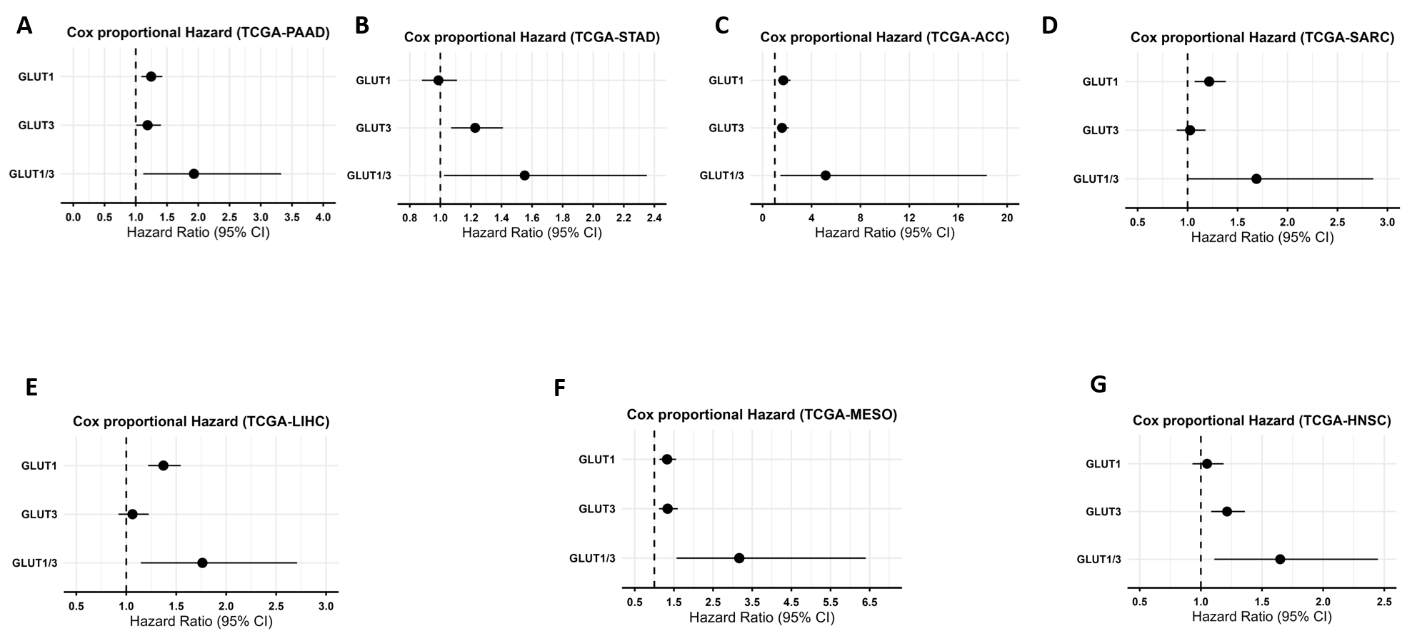

**Figure S6**. **Multivariate cox hazard analyses of** **patients with GLUT1/3 high-high vs low-low based on mRNA expression in TCGA cancer types. Significant values are shown.**

**S6A**. Multivariate cox proportional hazard model suggested poorer survival in combined group in TCGA-PAAD. **S4B**. Multivariate cox proportional hazard model suggested poorer survival in combined group in TCGA-STAD. **S4C**. Multivariate cox proportional hazard model suggested poorer survival in combined group in TCGA-ACC. **S4D**. Multivariate cox proportional hazard model suggested poorer survival in combined group in TCGA-SARC. **S4E**. Multivariate cox proportional hazard model suggested poorer survival in combined group in TCGA-LIHC. **S2F**. Multivariate cox proportional hazard model suggested poorer survival in combined group in TCGA-MESO. **S4G**. Multivariate cox proportional hazard model suggested poorer survival in combined group in TCGA-HNSC.

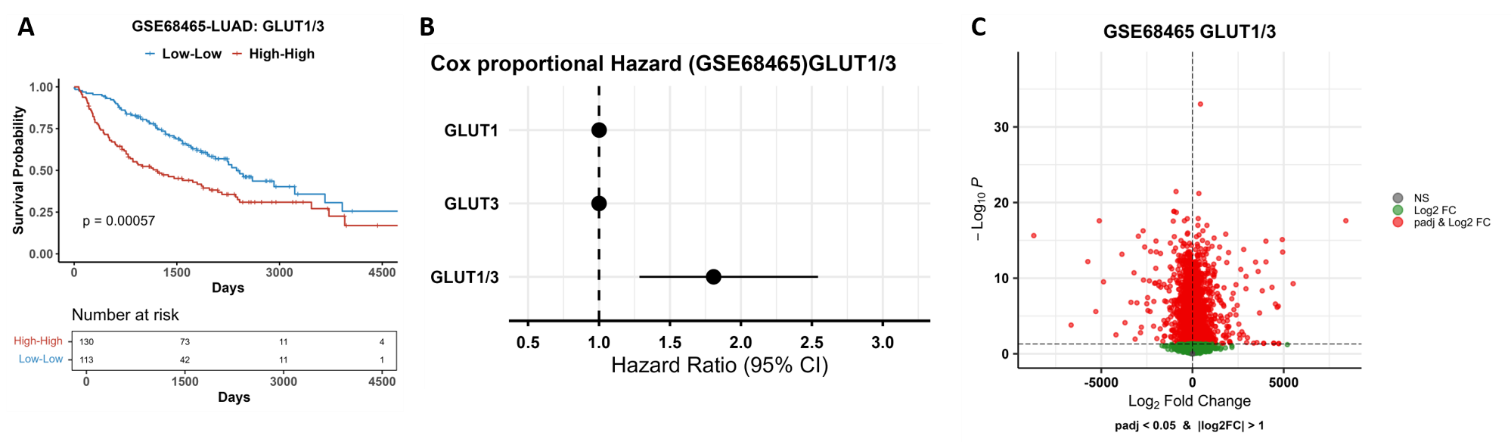

**Figure S7**. **GSE68465 LUAD cohort survival and volcano plot based on DEGs from GLUT1/3 high-high vs low-low groups**

**S7A**. Kaplan-Meier survival curve of GLUT1/3 shows that high-high patients have poorer survival vs low-low patients (P=0.00091) LUAD cohort. **S7B**. Multivariate cox proportional hazard model suggested poorer survival in combined group in LUAD.

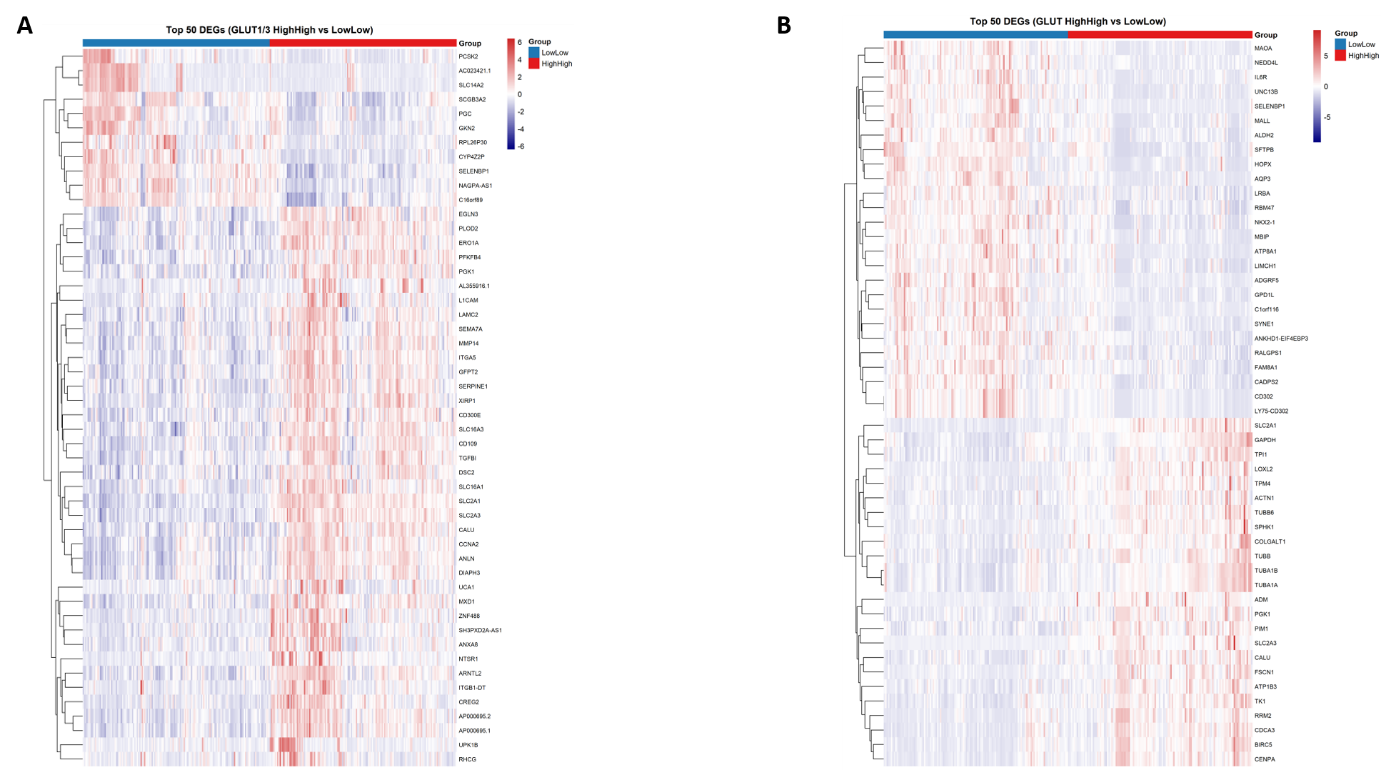

**Figure S8.** **Heatmap of top 50 DEGs in TCGA-LUAD and GSE68465 based on GLUT1/3 combination high-high vs low-low patients**

**S8A**. Heatmap of top 50 DEGs in TCGA patients based on GLUT1/3 high-high vs low-low. **S8B**. Heatmap of top 50 DEGs in GSE68465 patients based on GLUT1/3 high-high vs low-low. For all the analysis, FDR<0.5; logpadj<0.05 were used.

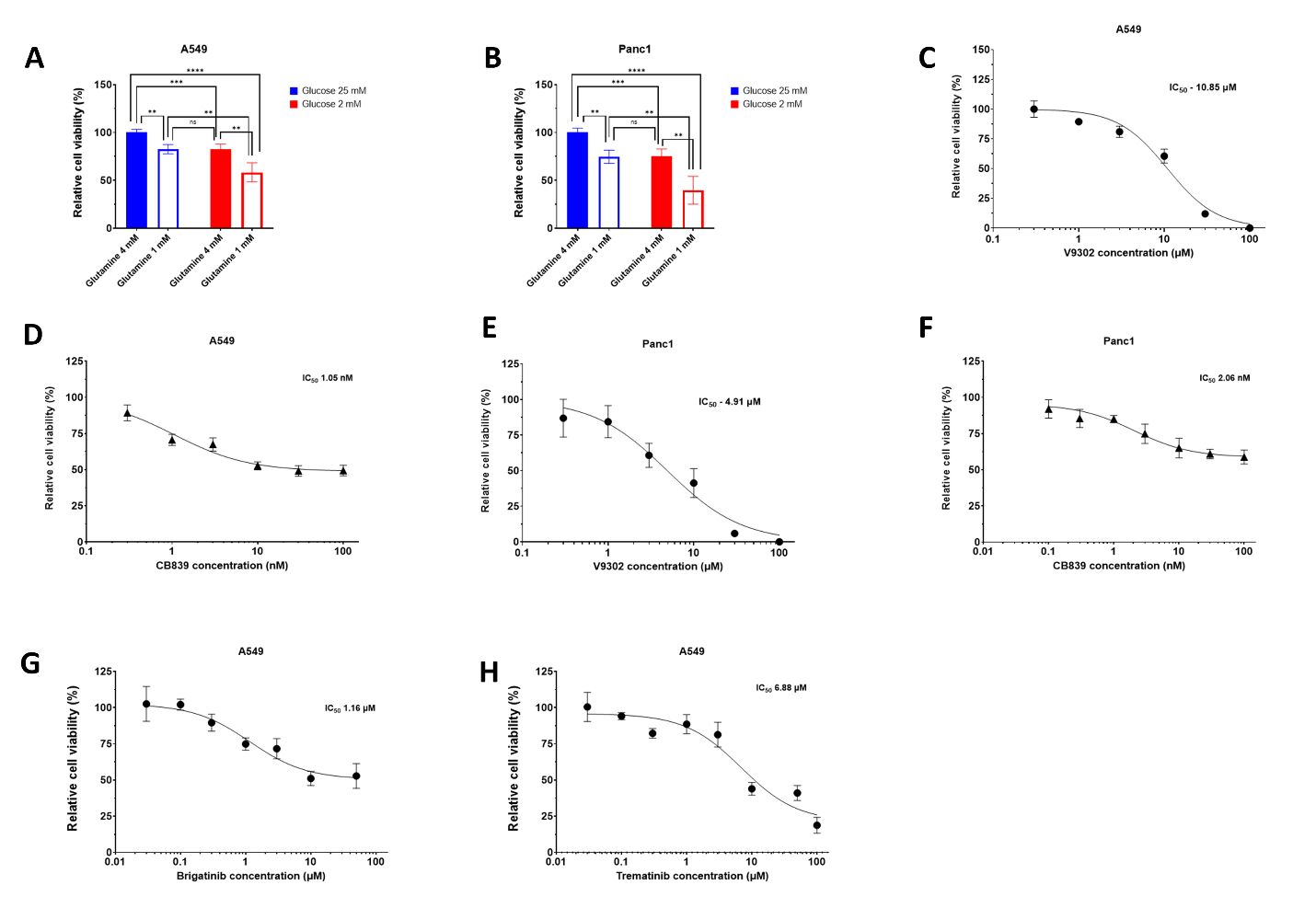

**Figure S9**. **Dose-dependent IC_50_ analysis of different pre-clinical or clinical anticancer agents in A549 and Panc1 cells.**

A549 or Panc1 cells were treated under different conditions for 48 hours, and then measured for their proliferation rates and their IC_50_.

**S9A.** Low glucose (2 mM) and low glutamine (1 mM) were significantly more toxic against A549 cells in comparison to high glucose (25 mM) and high glutamine (4 mM). **S9B.** Low glucose (2 mM) and low glutamine (1 mM) were significantly more toxic against Panc1 cells in comparison to high glucose (25 mM) and high glutamine (4 mM). **S9C**. Dose-dependent cell growth inhibition by glutamine transporter inhibitor V9302 in A549 cells. **S9D**. Dose-dependent cell proliferation inhibition by glutaminase inhibitor CB839 in A549 cells. **S9E**. Dose-dependent cell proliferation inhibition by glutamine transporter inhibitor V9302 in Panc1 cells. **S9F**. Dose-dependent cell proliferation inhibition by glutaminase inhibitor CB839 in Panc1 cells. **S9G**. Dose-dependent cell proliferation inhibition by ALK inhibitor brigatinib in A549 cells. **S9H**. Dose-dependent cell proliferation inhibition by MEK inhibitor trametinib in A549 cells.

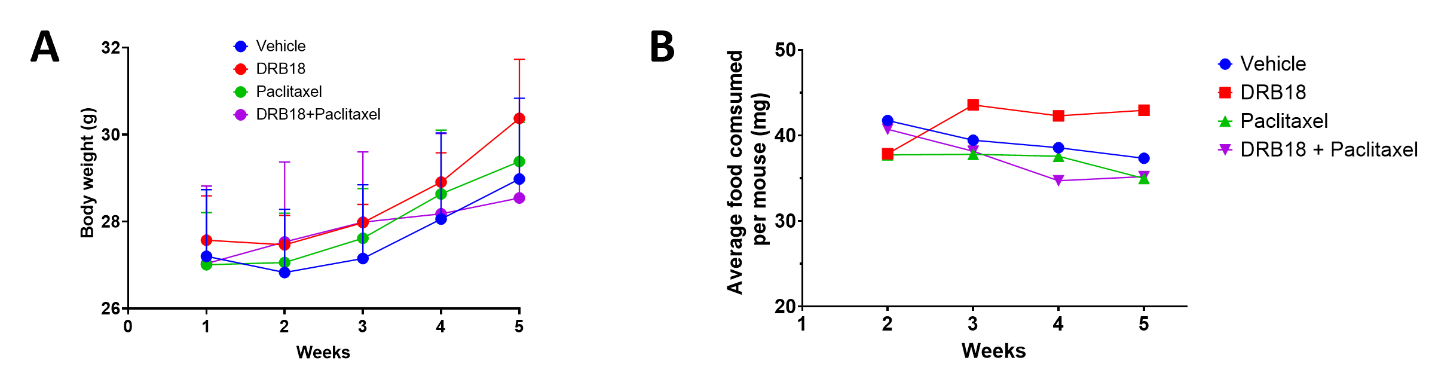

**Figure S10.** **Body weight and food consumption in tumor-bearing nude mice during 5 weeks of treatment.**

**S10A**. Body weight of mice after 5 weeks of treatment with vehicle, DRB18 alone, paclitaxel alone or DRB18 and paclitaxel combined. **S10B**. Food consumption per mouse during 5 weeks of treatment with vehicle, DRB18 alone, paclitaxel alone or DRB18 and paclitaxel combined.

**Supplementary Table 1. Clinical characteristics of LUAD patients in LC1531 Tumor microarray (TMA) used for GLUT1 analysis**

| Position | Age | Sex | Pathology diagnosis | TNM | Stage | Type | Tissue ID. |
| --- | --- | --- | --- | --- | --- | --- | --- |
| A1 | 69 | F | Adenocarcinoma | T2aN1M0 | IIB | Malignant | Rln020431 |
| A2 | 69 | F | Adenocarcinoma (trachea) | T2aN1M0 | IIB | Malignant | Rln020431 |
| A3 | 69 | F | Cancer adjacent lung tissue | - | - | AT | Rln020431 |
| A4 | 72 | M | Papillary adenocarcinoma | T1aN0M0 | IA | Malignant | Rln010183 |
| A5 | 72 | M | Papillary adenocarcinoma | T1aN0M0 | IA | Malignant | Rln010183 |
| A6 | 72 | M | Cancer adjacent lung tissue | - | - | AT | Rln010183 |
| B1 | 53 | M | Adenocarcinoma | T3N0M0 | IIB | Malignant | Rln030340 |
| B2 | 53 | M | Adenocarcinoma | T3N0M0 | IIB | Malignant | Rln030340 |
| B3 | 53 | M | Normal adjacent lung tissue | - | - | NAT | Rln030340 |
| C1 | 75 | M | Adenocarcinoma (cartilage tissue) | T2N0M0 | IB | Malignant | 37 |
| C2 | 75 | M | Adenocarcinoma | T2N0M0 | IB | Malignant | Rln020687 |
| C3 | 75 | M | Normal adjacent lung tissue | - | - | NAT | Rln020687 |
| C4 | 56 | M | Adenocarcinoma | T2aN0M0 | IB | Malignant | Rln030280 |
| C5 | 56 | M | Adenocarcinoma | T2aN0M0 | IB | Malignant | Rln030280 |
| C6 | 56 | M | Cancer adjacent lung tissue | - | - | AT | Rln030280 |
| D1 | 37 | F | Adenocarcinoma | T2bN0M0 | IIA | Malignant | Rln020019 |
| D2 | 37 | F | Adenocarcinoma | T2bN0M0 | IIA | Malignant | Rln020019 |
| D3 | 37 | F | Normal adjacent lung tissue | - | - | NAT | Rln020019 |
| D4 | 40 | M | Adenocarcinoma | T2aN0M0 | IB | Malignant | Rln020182 |
| D5 | 40 | M | Adenocarcinoma | T2aN0M0 | IB | Malignant | Rln020182 |
| D6 | 40 | M | Normal adjacent lung tissue | - | - | NAT | Rln020182 |
| E1 | 48 | F | Adenocarcinoma | T2bN0M0 | IIA | Malignant | Rln060003 |
| E2 | 48 | F | Adenocarcinoma | T2bN0M0 | IIA | Malignant | Rln060003 |
| E3 | 48 | F | Cancer adjacent lung tissue | - | - | AT | Rln060003 |
| E4 | 59 | F | Adenocarcinoma | T2aN1M0 | IIB | Malignant | Rln010077 |
| E5 | 59 | F | Adenocarcinoma | T2aN1M0 | IIB | Malignant | Rln010077 |
| E6 | 59 | F | Cancer adjacent lung tissue | - | - | AT | Rln010077 |
| G1 | 42 | F | Adenocarcinoma | T2aN0M0 | IB | Malignant | Rln020205 |
| G2 | 42 | F | Adenocarcinoma | T2aN0M0 | IB | Malignant | Rln020205 |
| G3 | 42 | F | Normal adjacent lung tissue | - | - | NAT | Rln020205 |
| G4 | 63 | F | Adenocarcinoma | T2bN0M0 | IIA | Malignant | Rln030394 |
| G5 | 63 | F | Adenocarcinoma | T2bN0M0 | IIA | Malignant | Rln030394 |
| G6 | 63 | F | Normal adjacent lung tissue | - | - | NAT | Rln030394 |
| H1 | 32 | F | Adenocarcinoma | T3N1M0 | IIIA | Malignant | Rln030245 |
| H2 | 32 | F | Adenocarcinoma | T3N1M0 | IIIA | Malignant | Rln030245 |
| H3 | 32 | F | Normal adjacent lung tissue | - | - | NAT | Rln030245 |
| H4 | 51 | F | Adenocarcinoma | T2aN0M0 | IB | Malignant | Rln030199 |
| H5 | 51 | F | Adenocarcinoma | T2aN0M0 | IB | Malignant | Rln030199 |
| H6 | 51 | F | Normal adjacent lung tissue | - | - | NAT | Rln030199 |
| H10 | 53 | M | Adenocarcinoma | T4N0M0 | IIIA | Malignant | Rln020521 |
| H11 | 53 | M | Adenocarcinoma | T4N0M0 | IIIA | Malignant | Rln020521 |
| H12 | 53 | M | Cancer adjacent lung tissue | - | - | AT | Rln020521 |
| H13 | 51 | M | Adenocarcinoma | T2aN0M0 | IB | Malignant | Rln010117 |
| H14 | 51 | M | Adenocarcinoma | T2aN0M0 | IB | Malignant | Rln010117 |
| H15 | 51 | M | Cancer adjacent lung tissue | - | - | AT | Rln010117 |
| I1 | 55 | M | Adenocarcinoma | T2aN0M0 | IB | Malignant | Rln020228 |
| I2 | 55 | M | Adenocarcinoma | T2aN0M0 | IB | Malignant | Rln020228 |
| I3 | 55 | M | Cancer adjacent lung tissue | - | - | AT | Rln020228 |
| I4 | 54 | F | Adenocarcinoma | T4N1M0 | IIIA | Malignant | Rln020020 |
| I5 | 54 | F | Adenocarcinoma | T4N1M0 | IIIA | Malignant | Rln020020 |
| I6 | 54 | F | Cancer adjacent lung tissue | - | - | AT | Rln020020 |
| J1 | 69 | M | Adenocarcinoma | T2bN0M0 | IIA | Malignant | Rln010090 |
| J2 | 69 | M | Adenocarcinoma | T2bN0M0 | IIA | Malignant | 53 |
| J3 | 69 | M | Normal adjacent lung tissue | - | - | NAT | Rln010090 |
| J4 | 64 | M | Adenocarcinoma | T1bN0M0 | IA2 | Malignant | Rln020238 |
| J5 | 64 | M | Adenocarcinoma | T1bN0M0 | IA3 | Malignant | Rln020238 |
| J6 | 64 | M | Normal adjacent lung tissue | - | - | NAT | Rln020238 |
| J13 | 66 | F | Adenocarcinoma | T2bN1M0 | IIB | Malignant | Rln030084 |
| J14 | 66 | F | Adenocarcinoma | T2bN1M0 | IIB | Malignant | Rln030084 |
| J15 | 66 | F | Normal adjacent lung tissue | - | - | NAT | Rln030084 |
| K1 | 59 | M | Adenocarcinoma | T3N1M0 | IIIA | Malignant | Rln010094 |
| K2 | 59 | M | Adenocarcinoma | T3N1M0 | IIIA | Malignant | Rln010094 |
| K3 | 59 | M | Cancer adjacent lung tissue | - | - | AT | Rln010094 |

AT- Cancer adjacent lung tissue, NAT – Normal adjacent lung tissue

**Supplementary Table 2. Clinical characteristics of LUAD patients in LC486 Tumor microarray (TMA) used for GLUT3 analysis**

| Position | Age | Sex | Pathology diagnosis | TNM | Stage | Type | Tissue ID. |
| --- | --- | --- | --- | --- | --- | --- | --- |
| A1 | 42 | M | Adenocarcinoma | T2N0M0 | I | Malignant | Rln010049 |
| B1 | 42 | M | Adenocarcinoma | T2N0M0 | I | Malignant | Rln010049 |
| C1 | 42 | M | Cancer adjacent lung tissue | - | - | adjacent | Rln010049 |
| A6 | 53 | F | Adenocarcinoma | T2N1M0 | II | Malignant | Rln010055 |
| B6 | 53 | F | Adenocarcinoma | T2N1M0 | II | Malignant | Rln010055 |
| C6 | 53 | F | Cancer adjacent lung tissue | - | - | adjacent | Rln010055 |
| A8 | 75 | M | Adenocarcinoma | T2N0M0 | I | Malignant | Rln010069 |
| B8 | 75 | M | Adenocarcinoma | T2N0M0 | I | Malignant | Rln010069 |
| C8 | 75 | M | Cancer adjacent lung tissue | - | - | adjacent | Rln010069 |
| A3 | 59 | F | Adenocarcinoma | T2N1M0 | II | Malignant | Rln010077 |
| B3 | 59 | F | Adenocarcinoma | T2N1M0 | II | Malignant | Rln010077 |
| C3 | 59 | F | Cancer adjacent lung tissue | - | - | adjacent | Rln010077 |
| A2 | 62 | M | Adenocarcinoma | T2N1M1 | IV | Malignant | Rln010078 |
| B2 | 62 | M | Adenocarcinoma | T2N1M1 | IV | Malignant | Rln010078 |
| C2 | 62 | M | Cancer adjacent lung tissue | - | - | adjacent | Rln010078 |
| A5 | 54 | M | Adenocarcinoma | T2N0M0 | I | Malignant | Rln010089 |
| B5 | 54 | M | Adenocarcinoma | T2N0M0 | I | Malignant | Rln010089 |
| C5 | 54 | M | Cancer adjacent lung tissue | - | - | adjacent | Rln010089 |
| A7 | 69 | F | Adenocarcinoma | T3N1M0 | IIIA | Malignant | Rln010181 |
| B7 | 69 | F | Adenocarcinoma | T3N1M0 | IIIA | Malignant | Rln010181 |
| C7 | 69 | F | Cancer adjacent lung tissue | - | - | adjacent | Rln010181 |
| A4 | 65 | F | Adenocarcinoma | T2N0M0 | I | Malignant | Rln020050 |
| B4 | 65 | F | Adenocarcinoma | T2N0M0 | I | Malignant | Rln020050 |
| C4 | 65 | F | Cancer adjacent lung tissue | - | - | adjacent | Rln020050 |

TNM -  T (Tumor size/invasion), N (Nodal status), and M (Metastasis)

**Supplementary Table 3. Clinical characteristics of LUAD patient in HuCAT232 Tumor microarray (TMA) used for GLUT1 analysis**

| Position | Age | Sex | Pathology diagnosis | TNM | Stage | Type |
| --- | --- | --- | --- | --- | --- | --- |
| A1 | 36 | M | Adenocarcinoma, stage IV | NA | NA | Malignant |

NA – Not available

**Supplementary Table 4. Clinical characteristics of patient in HuFPT178 Tumor microarray (TMA) used for GLUT1 analysis**

| Position | Age | Sex | Pathology diagnosis | TNM | Stage | Type | Tissue ID. |
| --- | --- | --- | --- | --- | --- | --- | --- |
| NA | NA | NA | Normal lung tissue | NA | NA | Normal Lung Tissue | NA |

NA – Not available

**Supplementary Table 5. Physiochemical properties, pharmacokinetics, drug-likeness, and medicinal chemistry for DRB18**

| **Physicochemical properties of DRB18** | | | **Recommended value** |
| --- | --- | --- | --- |
| Formula | | C_21_H_21_ClN_2_O_2_ |  |
| Molecular Weight | | 368.86 gm/mol |  |
| Number of Heavy atoms | | 26 |  |
| Number of aromatic atoms | | 18 |  |
| Fraction Csp^3^ | | 0.14 | ≤1 |
| Number of rotatable bonds | | 6 | ≤10 |
| Number of H-bond acceptors | | 2 | ≤12 |
| Number of H-bond donors | | 4 | ≤5 |
| Molar refractivity | | 108.05 |  |
| TPSA | | 64.52 Å^2^ | ≤140 Å^2^ |
| GI absorption | | High |  |
| **Drug-likeness** | | | |
| Lipinski | | Yes, O violation |  |
| Ghose | | Yes |  |
| Veber | | Yes |  |
| Egan | | Yes |  |
| Muegge | | Yes |  |
| Bioavailability score | | 0.55 (55%) |  |
| Synthetic Accessibility | | 2.34 | 1 (easy to make) and 10 (difficult to make) |
